# Revealing unseen dynamical regimes of ecosystems from population time-series data

**DOI:** 10.1101/2024.08.07.607005

**Authors:** Lucas P. Medeiros, Darian K. Sorenson, Bethany J. Johnson, Eric P. Palkovacs, Stephan B. Munch

## Abstract

Many ecosystems can exist in alternative dynamical regimes for which small changes in an environmental driver can cause sudden jumps between regimes. However, predicting the dynamics of regimes that occur under unobserved levels of the environmental driver has remained an unsolved challenge in ecology with important implications for conservation and management. Here we show that integrating population time-series data and information on the putative driver into an empirical dynamic model allows us to predict new dynamical regimes without the need to specify a population dynamics model. As a proof of concept, we demonstrate that we can accurately predict fixed-point, cyclic, or chaotic dynamics under unseen driver levels for a range of simulated models. For a model with an abrupt population collapse, we show that our approach can anticipate the regime that follows the tipping point. We then apply our approach to data from an experimental microbial ecosystem and from a lake planktonic ecosystem. We find that we can reconstruct transitions away from chaos in the experimental ecosystem and anticipate the dynamics of the oligotrophic regime in the lake ecosystem. These results lay the groundwork for making rational decisions about preventing, or preparing for, regime shifts in natural ecosystems.

## Introduction

Many ecosystems can exist in several distinct alternative dynamical regimes, and rapid shifts from one regime to another may result from gradual changes in some environmental driver or “control parameter”.^1, 2^ Well-documented examples of regime shifts include clear macrophyte-dominated lakes transitioning into turbid phytoplankton-dominated lakes due to nutrient input,^3^ kelp forests becoming sea urchin barrens due to predator loss,^4^ coral reefs giving way to macroalgae reefs as a result of warming,^5^ and forests shifting to savannas after changes in precipitation.^6^ Because ecosystem services depend greatly on the state of the system, there has been considerable interest in understanding and predicting the dynamical regimes that may emerge under changing environmental conditions.

Predicting ecological regime shifts in time to avert them has been an important avenue of research and many Early Warning Signals (EWS) have been developed to do so.^7^ These signals can be computed directly from population time-series data and include variance, skewness, the AR(1) coefficient, and the power spectrum.^8^ The unifying idea behind EWS is that changes in the environmental driver cause the ecosystem to cross a tipping point (i.e., a bifurcation) and most are based on “critical slowing down”—in which the stability of a fixed point relaxes as the bifurcation is approached.^9^ Empirical evidence for critical slowing down has been found in plankton in lakes,^10^ photo-inhibition in cyanobacterial populations,^11^ declining food sources for *Daphnia magna*,^12^ and increased mortality in yeast.^13^ A key feature of EWS is that they are generic, that is, the dynamical behavior close to a bifurcation is well-described by a handful of canonical “normal forms”^14^ such that we do not need detailed knowledge of the underlying dynamics to anticipate a shift. However, EWS pertain to a relatively narrow range of bifurcations—namely the fold, transcritical, and Hopf bifurcations^2, 15^— and approaches that apply to a wider range of cases are currently needed. Most importantly, EWS only tell us that the ecosystem might be approaching a regime shift but give no information about what the upcoming dynamics will look like.

Predicting the dynamics of ecosystems under unobserved values of the environmental driver, such as beyond a proposed tipping point, has been a major challenge in ecology. For example, anticipating not only the structure but the population dynamics of planktonic food webs under nutrient levels not yet observed is notoriously difficult.^16^ The difficulty stems from accurately extrapolating the effect of the driver on species abundances using data on few species and over a limited range of the driver. Unfortunately, most EWS are unable to perform this extrapolation^17, 18^ and the few approaches that do (e.g., approaches leveraging deep learning^19^) require prior knowledge of the class of bifurcations. Although model-based approaches provide more specific predictions,^2, 20^ these require considerably more detailed knowledge about the underlying dynamics than generic EWS and are unlikely to be transferable across ecosystems. Therefore, an alternative approach, with generality similar to that of existing EWS, but that also predicts the dynamics beyond a tipping point is highly desirable.

Empirical Dynamic Modeling (EDM) is a framework based on nonparametric function approximation and time-delay embedding that has proved useful to understand and forecast a wide range of dynamical systems. Examples include forecasting recruitment in fisheries,^21^ understanding the environmental drivers of diseases,^22^ inferring causal interaction networks,^23^ and quantifying population responses to perturbations.^24^ By learning about system dynamics directly from time-series data (through nonparametric function approximation) and by accounting for unobserved state variables (through time-delay embedding), EDM is able to give insights about the dynamics without the need to specify a model, as with EWS. Several studies have used EDM to develop new EWS, which include approaches based on prediction accuracy,^25^ nonlinearity,^26^ and eigenvalues of the Jacobian matrix.^27^ Other recent studies have used EDM to detect if and when a regime shift has happened in the past based on prediction accuracy tests.^28, 29^ What is currently lacking from both EWS and EDM approaches is the capacity to predict dynamics beyond observed driver levels. Yet, being able to leverage information on putative drivers of regime shifts (e.g., temperature, nutrients, rainfall, fishing pressure) could allow us to anticipate unseen regimes and at the same time provide some mechanistic understanding.

Here we introduce a novel approach based on EDM that allows us to predict the dynamical behavior of an ecosystem at unobserved levels of an environmental driver. Our approach works by first integrating population time-series data and information on the putative driver into a Gaussian Process regression with time-delay embedding (GPEDM,^30^) to learn how the driver affects population dynamics. We then extrapolate the dynamics to unobserved levels of the driver to predict unseen dynamical regimes. Using several population dynamics models, we demonstrate that we can accurately predict unseen fixed-point, cyclic, or chaotic regimes, in addition to predicting the location of multiple tipping points in a bifurcation diagram. We also show that, for a model with an abrupt population collapse, our approach can anticipate the regime that follows the tipping point. Finally, we apply our approach to empirical time series from both an experimental microbial ecosystem and a lake planktonic ecosystem. We show that we can reconstruct transitions away from chaos in the experimental ecosystem and anticipate the dynamics of the oligotrophic regime in the lake ecosystem.

## Results

### Testing approach on model-generated data

We first tested the ability of our approach to predict unseen dynamical regimes and the location of tipping points using four discrete-time population dynamics models (see *Methods*). For each model, we trained a GP-EDM model (*Supplementary Note 1* ;^30^) using time series from a single species (*x*_*i*_) generated at four different levels of an environmental driver (i.e., a control parameter *p*; Fig. 1a). The trained GP-EDM models produced highly accurate leave-one-out predictions under the observed data (*R*^2^: single-species model, 0.992; two-species competition model with harvesting, 0.998; two-species predatorprey model, 0.990; three-species competition model, 0.983; Fig. S1). Then, using the function *x*_*i*_(*t* + 1) = *G*_*i*_[*x*_*i*_(*t*), …, *x*_*i*_(*t* − *E*), *p*], approximated via GP-EDM (Fig. 1b), we performed predictions of *x*_*i*_ over unobserved values of *p* to reconstruct the bifurcation diagram of each model (Fig. 1c, d).

**Fig. 1.**
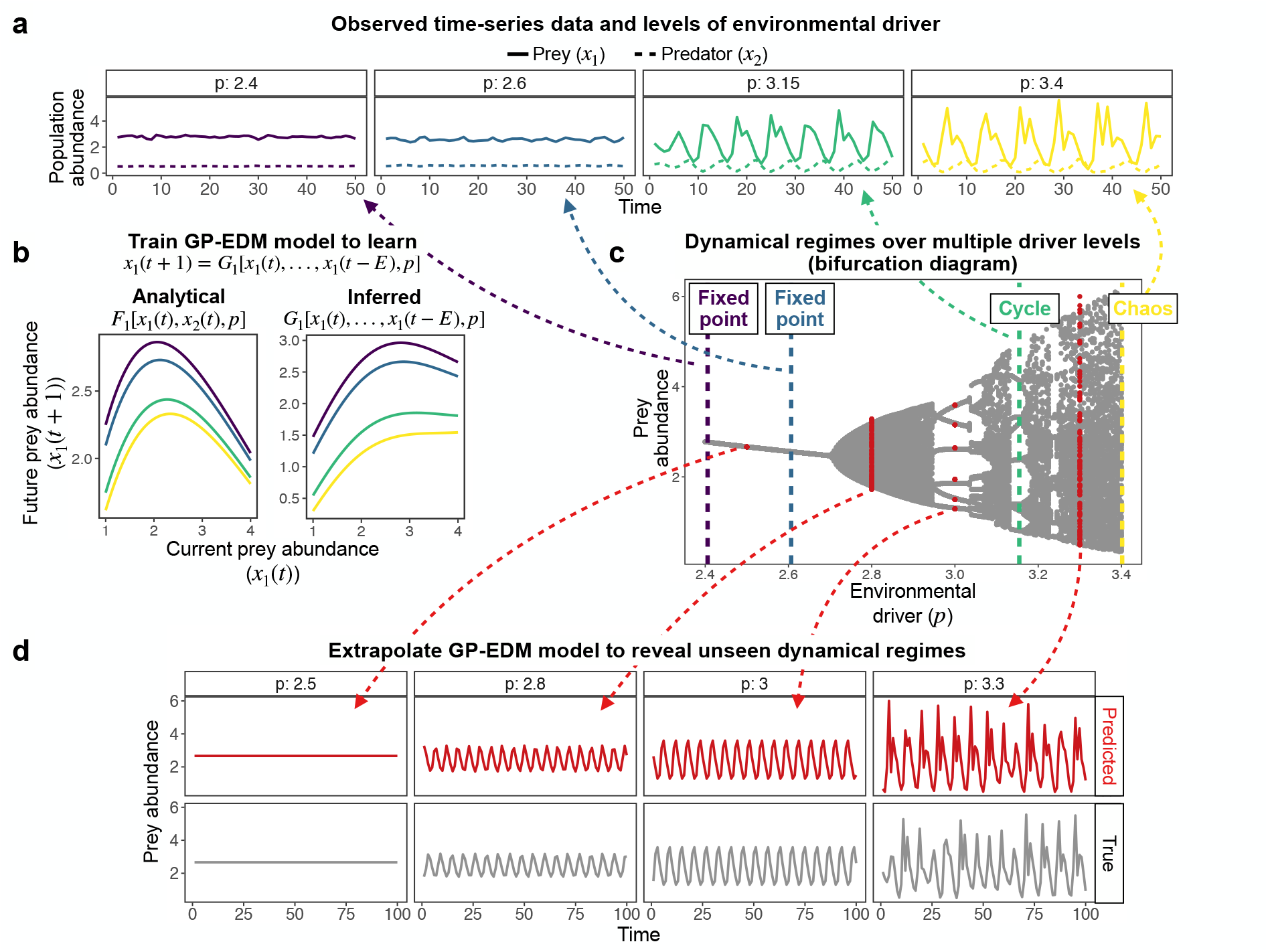
GP-EDM approach to reveal unseen dynamical regimes. **a**, Population time series generated from a predator-prey model (equation (9)) at four different levels of the attack rate parameter (i.e., the environmental driver *p*). Low values of *p* generate fixed points (*p* = 2.4 in purple and *p* = 2.6 in blue) whereas high values of *p* generate cycles (*p* = 3.15 in green) or chaos (*p* = 3.4 in yellow). **b**, Left panel depicts the true relationship between current (*x*_1_(*t*)) and future (*x*_1_(*t* + 1)) prey abundance at different levels of *p* (different colors). Right panel depicts the same relationship but obtained from the trained GP-EDM model. **c**, Bifurcation diagram of the predator-prey model showing the different dynamical regimes that emerge as we vary *p* (gray points). The four regimes shown in **a** are represented as vertical dashed lines. We can then extrapolate our trained GP-EDM to unseen levels of *p* that are shown in red. **d**, The predicted dynamical regimes at these unseen levels of *p* (in red) match closely the true dynamics (in gray). These dynamical regimes are: fixed point (*p* = 2.5), low-amplitude cycle (*p* = 2.8), high-amplitude cycle (*p* = 3.0), and chaos (*p* = 3.3).

We were able to accurately predict the dynamics of *x*_*i*_ across many unobserved levels of *p* for all four models (Fig. 2). For instance, having observed the prey time series at four regimes of a 2-species predator-prey model (fixed points at *p* = 2.4 and *p* = 2.6, cycle at *p* = 3.15, and chaos at *p* = 3.4), we accurately predicted unobserved fixed-point, cyclic, or chaotic regimes (Fig. 2c). Predicted dynamical regimes significantly matched true regimes when measuring the similarity between them with the Jensen-Shannon divergence (average divergence: single-species model, 0.074; two-species competition model with harvesting, 0.124; two-species predator-prey model, 0.042; three-species competition model, 0.306). Note that our approach can still provide a qualitatively correct prediction even when the Jensen-Shannon divergence indicates a quantitatively wrong prediction (e.g., fixed point in Fig. 2d). As expected, our approach was most accurate when we used all species in the GP-EDM model (i.e., native coordinates) instead of using only a single species with time-delay embedding (i.e., delay coordinates; Fig. S2). Although with slightly less accuracy, we also detected the location of tipping points (i.e., bifurcations) that separate different dynamical regimes. This includes the location of period doubling bifurcations (Fig. 2a, d), a Hopf bifurcation (Fig. 2c), and a fold bifurcation associated with population collapse (Fig. 2b). We evaluated the sensitivity of our results by performing these analyses for different dynamical regimes in our training data (Fig. S3). We found that we can most accurately predict unseen regimes and detect the location of tipping points when the training data includes a chaotic regime, which provides information for the GP-EDM model across the entire range of *x*_*i*_ (i.e., across the entire state space). We also performed these analyses for several different training data sets with different amounts of process noise. Although high noise can make our approach less precise, we found that, on average, the Jensen-Shannon divergence between true and predicted dynamics was low (Fig. S4).

**Fig. 2.**
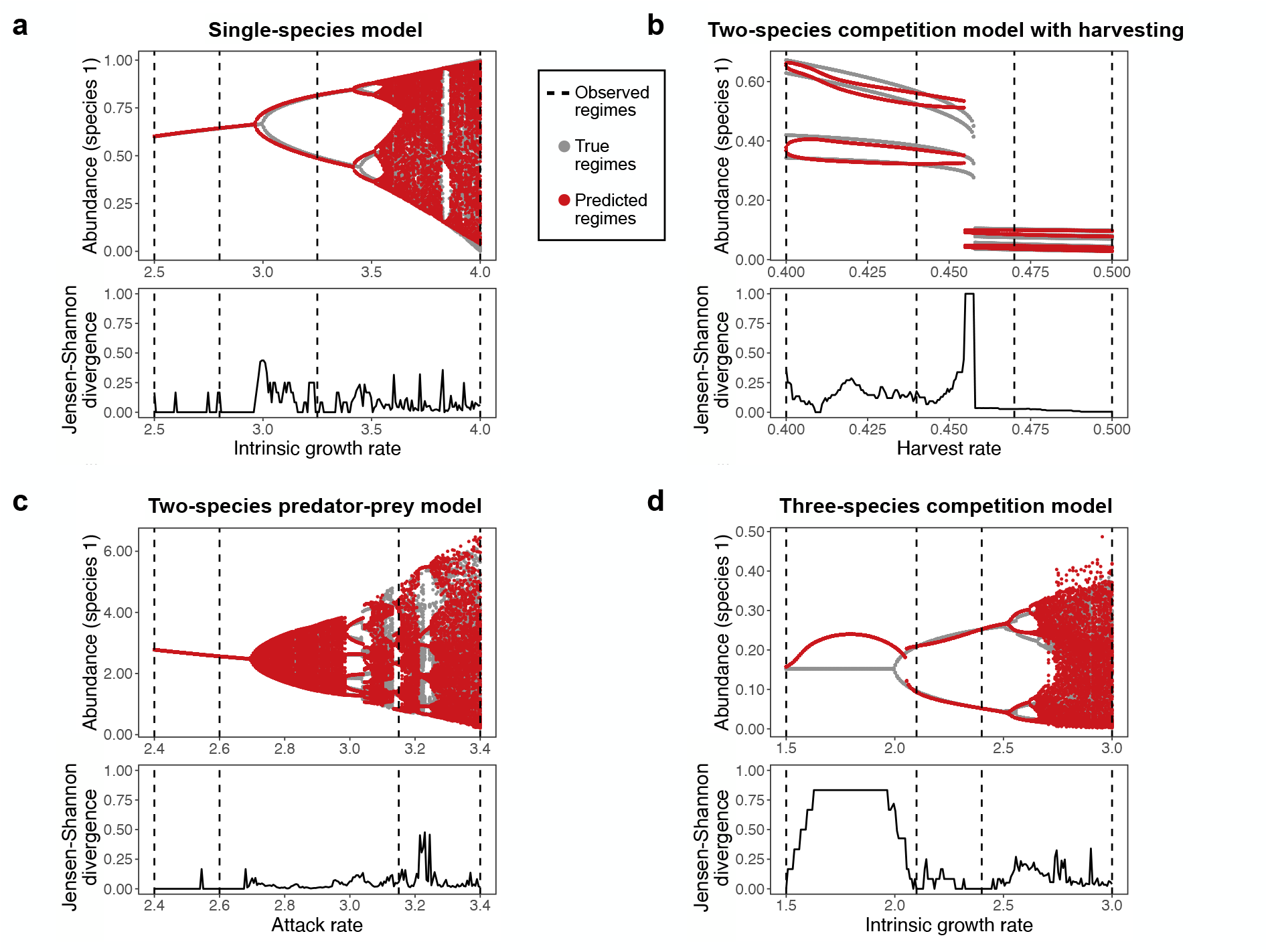
GP-EDM approach can accurately predict unseen dynamical regimes across a range of models. **a**-**d**, Top panels show the true (in gray) and predicted (in red) bifurcation diagrams for the four discrete-time population dynamics models (equations (7) to (10)). Bifurcation diagrams depict the population abundance of a given species after the transient period (y-axis) for a range of control parameter values (*p*, x-axis). Bottom panels show the Jensen-Shannon divergence computed at each value of *p*. Values closer to zero represent a closer match between the true and predicted dynamics. In all panels, the vertical dashed lines depict the values of *p* for which we observed population time series to train the GP-EDM model (as in Fig. 1c).

We then tested whether our approach can be used as an Early Warning Signal (EWS) of an upcoming population collapse using a 2-species competition model with harvesting (see *Methods*). To do so, we generated a time series for which the harvest rate (i.e., the control parameter *p*) changed linearly from 0.42 to 0.48 and a tipping point occurred at *p* = 0.464 according to a simple change-point analysis (Fig. 3a). We then sequentially updated our GP-EDM model as the training data set increased towards the tipping point and, for each update, we extrapolated the entire bifurcation diagram (i.e., all dynamical regimes from *p* = 0.42 to *p* = 0.48). Again, the best trained GP-EDM model produced highly accurate leave-one-out predictions (*R*^2^: 0.986).

**Fig. 3.**
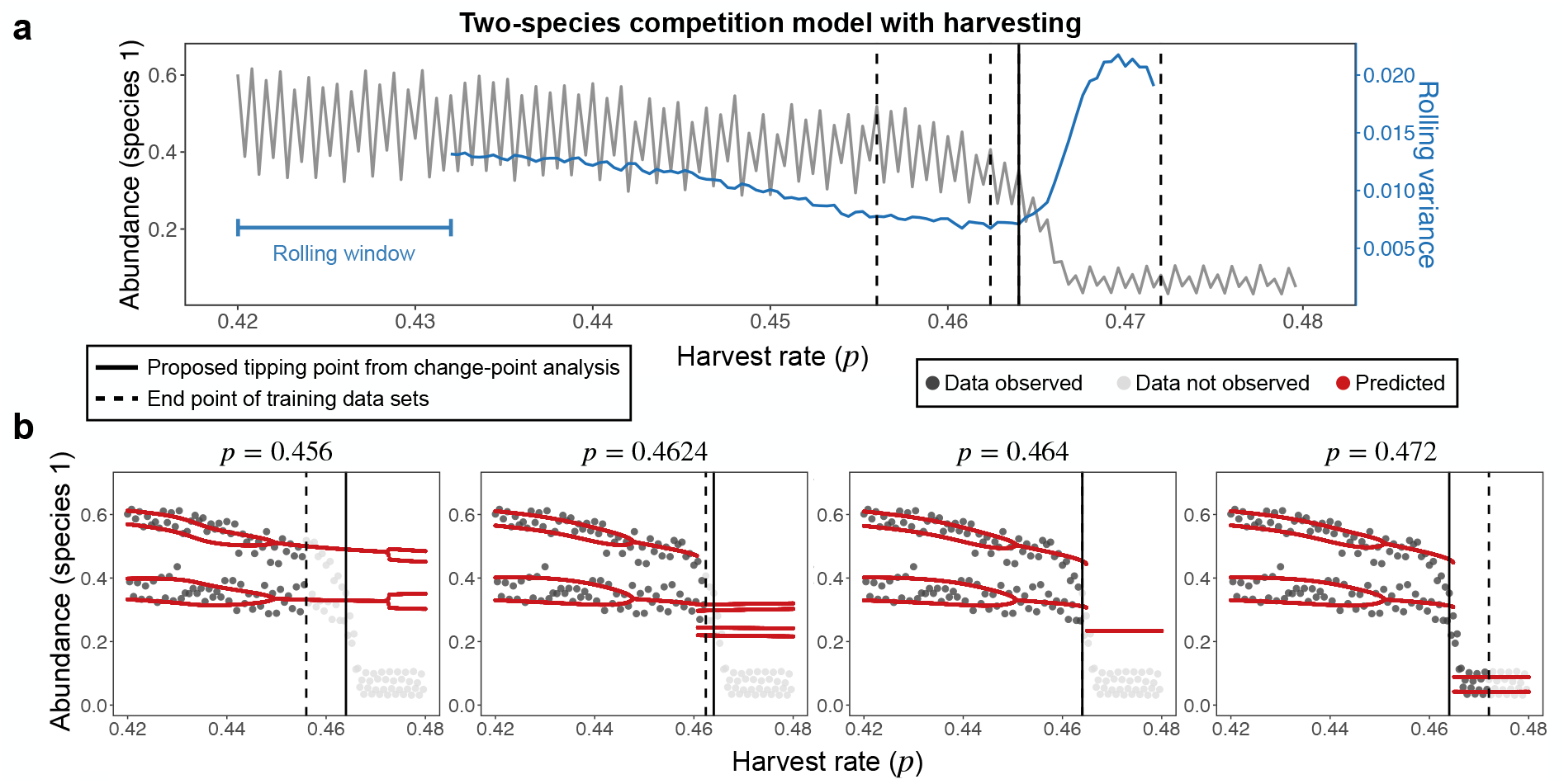
Illustration of GP-EDM approach as an Early Warning Signal. **a**, Population time series (in gray) of the harvested species in a two-species competition model with harvesting (equation (8)). Harvest rate (*p*) is depicted in the x-axis and is proportional to time. The population shows a tipping point at *p* = 0.464 (vertical solid line) according to a change-point analysis. The vertical dashed lines show the end points of the training data sets used with our GP-EDM model. The rolling variance (a typical EWS) computed using a window with 30 points is shown in blue. **b**, Predicted bifurcation diagrams (in red) using the GP-EDM model trained up to the end of each of the four time-series windows. Bifurcation diagrams depict the population abundance after the transient period (y-axis) for a range of harvest rate values (x-axis). The value of *p* at the end of the time series window is shown at the top of each plot. Gray points denote population abundance values used to train the GPEDM model (dark gray) or not yet observed (light gray).

Our approach was able to anticipate population collapse when the training data ended 4 time steps before the tipping point (Fig. 3b). Although the predicted abundance at the unseen regime was higher than the true abundance, our approach clearly anticipated an abrupt abundance shift. The harvest rate of population collapse was predicted to be *p* = 0.4608, which is slightly below the tipping point location (i.e., *p* = 0.464). When the training data ended exactly at the tipping point, our approach predicted a more accurate tipping point location at *p* = 0.465. Finally, when the training data contained the collapsed regime, our approach provided a very accurate reconstruction of the entire bifurcation diagram, as expected. The rolling variance of population abundance began to steadily increase at the tipping point (Fig. 3a), slightly after our approach first signalled a tipping point (Fig. 3b). This result suggests that these approaches can be used in concert to produce better EWS. When using training data with tipping points at different locations, we obtained similar although slightly less accurate predictions (Fig. S5).

### Applying approach to an experimental microbial ecosystem

Next, we applied our approach to population time-series data from a seminal chemostat experiment that discovered transitions from fixed point to chaos to cycles as dilution rate (*p*) was reduced.^31^ This data set is analogous to our simulation set up from Fig. 2 and, therefore, we used the same analytical approach (see *Methods*).

For each of the three species in the system, we obtained reasonable leave-one-out predictions for the observed abundances (*R*^2^: predator, 0.490; preferred prey, 0.508; less-preferred prey, 0.670; Fig. S6), suggesting that GP-EDM is correctly approximating the function *x*_*i*_(*t* + 1) = *G*_*i*_[*x*_*i*_(*t*), …, *x*_*i*_(*t* − *E*), *p*] at the observed dilution rates. More importantly, the bifurcation diagrams inferred independently from each species exhibit a large region of chaos around *p* = 0.5/day, cycles around *p* = 0.45/day and slightly above *p* = 0.6/day, and a fixed point for *p >* 0.7/day (Fig. 4). Our results also reveal the dilution rate levels at which we should observe unseen bifurcations. For example, the appearance of a fixed point is predicted to occur somewhere between *p* = 0.62/day and *p* = 0.68/day (Fig. 4). The fact that similar results were obtained independently for each species increases our confidence that we are accurately revealing previously unknown regimes and bifurcations. Results for predator and preferred prey species are robust to changes in GP-EDM implementation details, whereas results for the less-preferred prey species are more sensitive to these changes (Fig. S7).

**Fig. 4.**
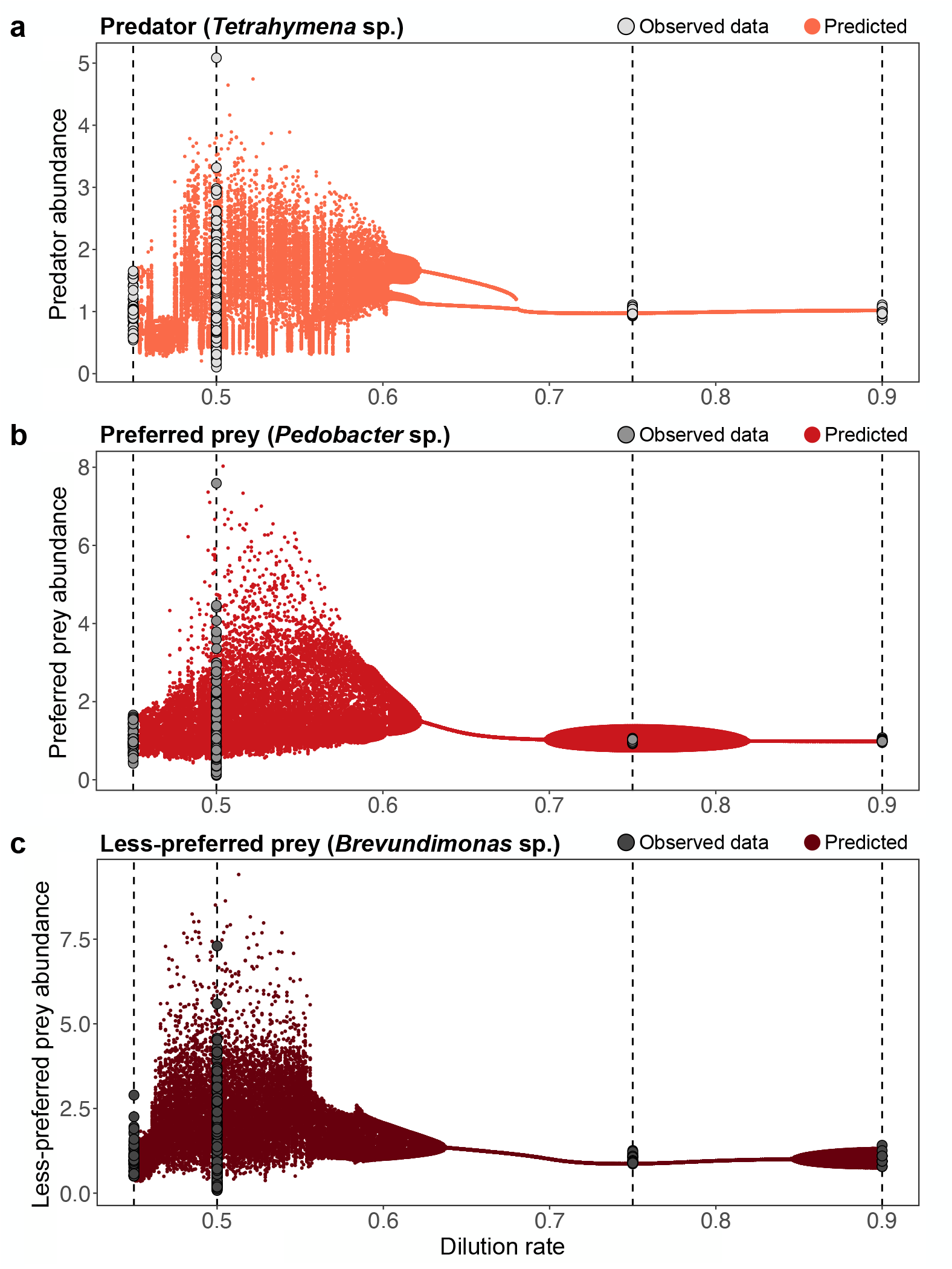
Predicting unseen dynamical regimes in an experimental microbial ecosystem. **a**-**c**, Bifurcation diagrams predicted with our GP-EDM approach (red points) for the predator species (*Tetrahymena* sp., light red, top panel), preferred prey species (*Pedobacter* sp., regular red, middle panel), and less-preferred prey species (*Brevundimonas* sp., dark red, bottom panel). Bifurcation diagrams depict the population abundance of a given species after the transient period (y-axis) for a range of dilution rate values (*p*, x-axis). Vertical dashed lines represent the dilution rates of the experimental treatments (0.45/day, 0.5/day, 0.75/day, and 0.9/day). Gray points show the abundance values at these dilution rates used to train a GP-EDM model separately for each species. Abundances are in arbitrary units due to our data scaling process (see *Methods*).

### Applying approach to a lake planktonic ecosystem

For our last set of analyses, we applied the EWS version of our approach to the wellknown case of lakes that shift from an eutrophic to an oligotrophic regime. We used monthly time-series data from Lake Zurich of 13 plankton functional groups, phosphate concentration (i.e., the control parameter *p*), and water temperature from January, 1978 to December 2019 (Fig. S8; *T* = 504 data points;^16^). Because this lake food web is thought to have shifted from an eutrophic to an oligotrophic regime via a gradual reduction of phosphate concentration,^16^ it is analogous to our simulations from Fig. 3.

We first performed a change-point analysis to determine the location of a potential tipping point for each functional group. We found that large green algae had the most well-resolved shift followed by small cryptophytes and omnivores (Table S1). This potential tipping point for large green algae occurred on November, 1987 (*t*_split_ = 119) at a phosphate concentration of 0.0472 mg*/*l (Fig. 5a). The potential tipping points for small cryptophytes (October, 1988; *t*_split_ = 130) and omnivores (June, 1986; *t*_split_ = 102) occurred around the same time, suggesting an ecosystem-wide shift.

**Fig. 5.**
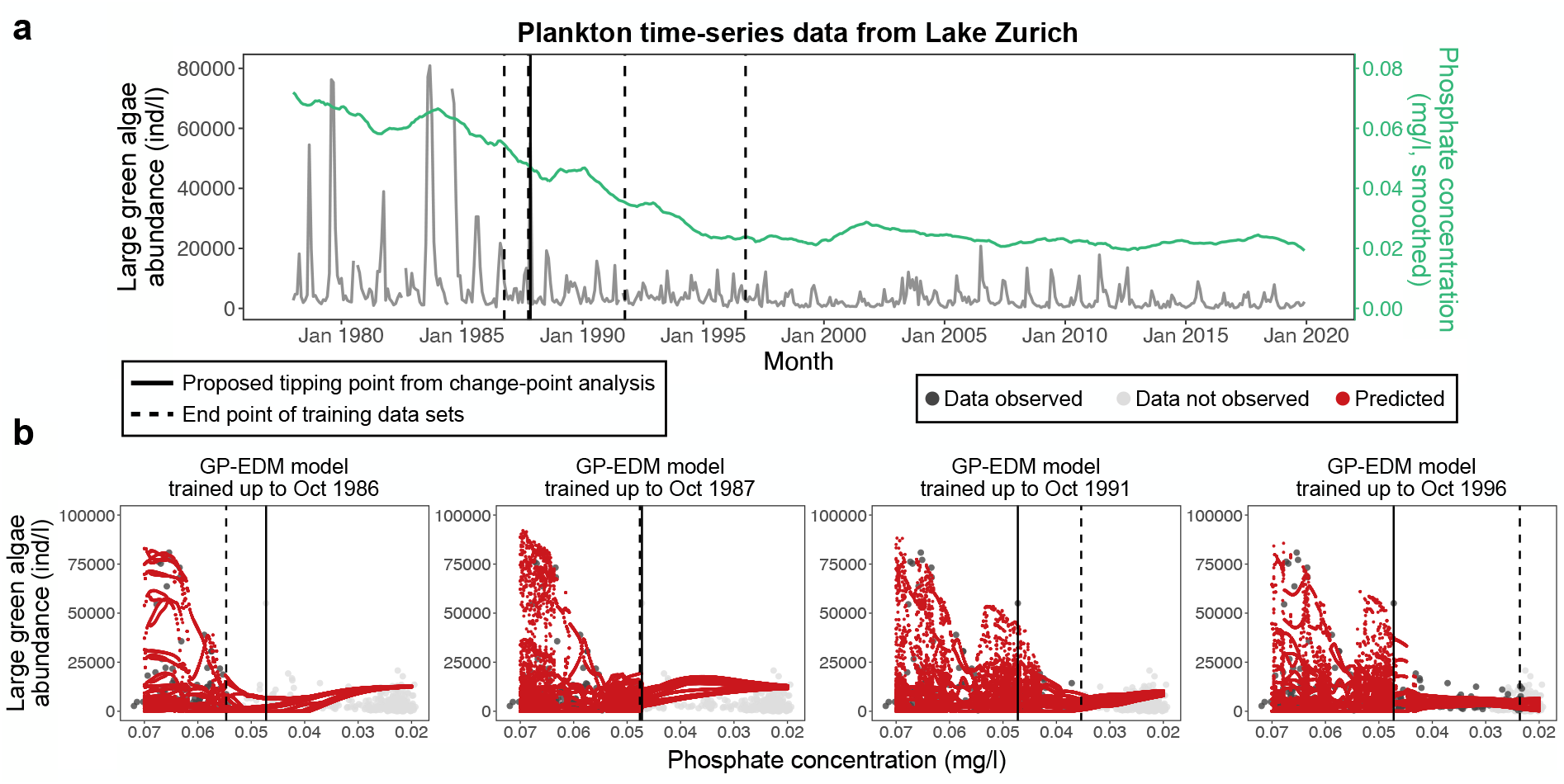
Anticipating the oligotrophic regime in a lake planktonic ecosystem. **a**, Monthly population time series (in gray) of large green algae in Lake Zurich from January, 1978 to December 2019. The population shows a tipping point on November, 1987 (vertical solid line) according to a change-point analysis. The vertical dashed lines show the end points of the training data sets used with our GP-EDM model. Phosphate concentration (smoothed via a centered moving average), a key environmental driver for re-oligotrophication in this system, is shown in green. **b**, Predicted bifurcation diagrams (in red) using the GP-EDM model trained up to the end of each of the four time-series windows. Bifurcation diagrams depict the population abundance after the transient period (y-axis) for a range of phosphate concentration values (x-axis). The time series end points are shown as dates at the top of each plot. Gray points denote population abundance values used to train the GP-EDM model (dark gray) or not yet observed (light gray).

To separate the effects of the control parameter from seasonality, we first applied a centered moving average smoother to the phosphate time series, removing seasonal oscillations while retaining the trend. Next, we included seasonality in our GP-EDM model by adding water temperature as an input. We then used data from January, 1978 to October, 1986 (106 data points) to train GP-EDM models and verify which functional groups were most predictable. Small green algae showed the best (*R*^2^ = 0.801), whereas large green algae showed the second-best (*R*^2^ = 0.770) leave-one-out prediction accuracy (Table S2). Due to strong evidence of a tipping point for large green algae from the change-point analysis, we focused subsequent analyses on this functional group.

We then reconstructed bifurcation diagrams for large green algae using successively larger training data sets (Fig. 5a). We found evidence of a previously unseen low abundance regime when the training data ended 1 year before the potential tipping point (i.e., training data up to October, 1986; Fig. 5b). We then clearly predicted an abrupt shift in the dynamics of large green algae when the training data ended 1 month before the potential tipping point (i.e., training data up to October, 1987; Fig. 5b). This abrupt shift was predicted to occur at a phosphate concentration of 0.0470 mg*/*l, very close to the phosphate concentration at the potential tipping point detected via change-point analysis. Interestingly, the bifurcation diagram reconstructed with training data up to October, 1987 was very similar to the bifurcation diagram reconstructed with training data up to October, 1996 (i.e., 9 years after the tipping point; Fig. 5b), confirming that the former predictions were accurate. These results were found to be robust to modest changes in GP-EDM implementation details (Fig. S9). In contrast to these results, the rolling variance of large green algae abundance—the well-known EWS that we used in Fig. 3—did not steadily increase prior to the tipping point (Fig. S10).

## Discussion

Foreseeing the dynamical regimes of ecosystems under unobserved environmental conditions is an important ecological challenge, with applications in both conservation and management.^1, 18, 32, 33^ Although a large body of research has developed Early Warning Signals (EWS) that foretell an impending regime shift,^7, 8^ anticipating what type of dynamics we should observe after the shift has remained elusive. The GP-EDM approach introduced here integrates population time-series data and information on a putative environmental driver to enable extrapolation to previously unseen dynamical regimes and serve as an EWS. In both simulated and empirical data, we found that GP-EDM can provide useful inference on when and how these systems change to a different regime.

Many EWS have been developed and employed to detect upcoming regime shifts, which include variance, skewness, the AR(1) coefficient, and the power spectrum.^8^ Although these indicators cannot give information about unseen dynamical regimes, other recent EWS based on deep learning or reservoir computing can perform this task.^19, 34, 35^ However, these approaches require large amounts of training data, information on the nonlinearities present in the underlying dynamics, or knowledge of the class of bifurcations. In contrast, GP-EDM can be applied with reasonable amounts of data and is completely agnostic about the underlying population dynamics. Therefore, our approach is able to identify bifurcations that are not among those typically considered in ecological studies. Most importantly, our approach leverages information on the putative driver of regime shifts and, by doing so, allows us to obtain more mechanistic insights. For instance, in Fig. 1b we inferred how density dependence changes with the control parameter *p* to cause bifurcations.

Although our results are promising, several limitations of GP-EDM are important to note. First, to be used as an EWS, the GP-EDM model must extrapolate to new regions of the control parameter space. Since the prediction accuracy of any Gaussian Process regression declines with long-range extrapolation,^36, 37^ this limits our ability to anticipate distant regime changes. Second, we found that when the training data includes a limited range of abundances (e.g., when it contains mostly fixed points), GP-EDM is likely to determine that the control parameter has little to no effect and the resulting bifurcation diagram is inaccurate (Fig. S3). This is unlikely to be an issue with natural population time series in which large deterministic fluctuations are common.^38, 39^ Third, ecosystems are typically under the effect of multiple environmental drivers. Our study, like most work in EWS, was restricted to a single control parameter that lead to codimension one bifurcations. Future studies need to evaluate the utility of GP-EDM and other EWS when multiple control parameters vary. Finally, in contrast to other EWS, GP-EDM requires data on the putative control parameter, or a suitable proxy. In the absence of a known driver, slow feature analysis^40^ may be useful to construct a suitable proxy and future studies could explore this idea.

In addition to testing our approach with model-generated data (Figs. 2 and 3), we applied our approach to two empirical data sets: an experimental microbial ecosystem (Fig. 4;^31^) and a lake planktonic ecosystem (Fig. 5;^16^). Both applications revealed unknown dynamical regimes and shifts between regimes in these ecosystems. Regarding the experimental microbial ecosystem, previous work had determined the existence of three qualitative distinct regimes (limit cycle at low dilution rate, chaos at intermediate dilution rate, and fixed point at high dilution rate;^31^). However, no experiment was conducted to verify the exact location of bifurcations. Our analyses shed light on this question, especially regarding the shift from the chaotic to the fixed point regime, which should occur at a dilution rate of 0.62-0.68/day (Fig. 4). Future experimental work could try to corroborate our prediction by performing experiments under these dilution rates.

Regarding the lake planktonic ecosystem, a large body of research has explored the transitions from oligotrophic to eutrophic and vice versa.^3, 16, 41, 42^ When analyzing a complex ecosystem such as a lake food web with multiple functional groups, it is challenging to determine whether a shift has occurred, because only certain groups may show a clear pattern of a tipping point.^18, 43–45^ In this sense, a key component of GP-EDM is that it uses time-delay embedding to account for hidden state variables. By focusing on large green algae—a highly predictable functional group that showed clear evidence of a tipping point—we were able to anticipate a possible ecosystem-wide regime shift. That is, one month before the potential tipping point, our approach clearly indicated an abrupt shift from a regime with large-amplitude abundance fluctuations to a regime with low-amplitude abundance fluctuations for large green algae. Note that this type of regime shift is common in phytoplankton under nutrient reductions, but is hard to anticipate with traditional EWS such as rolling variance (Fig. S10).^43, 45^ Here, we found that the shift from eutrophic to oligotrophic ocurred at a phosphate concentration of 0.0470 mg*/*l. An important area for future research is to apply GP-EDM to a range of natural ecosystems for which there is also data on a known environmental driver that induces regime shifts to obtain cross-system insights. This type of study could, for example, verify whether there is a typical phosphate concentration for the eutrophic-oligotrophic transition in lakes.

In summary, we show that GP-EDM accurately reveals previously unseen dynamical regimes under changing environmental conditions across a range of complexity. Although regime shifts can be extremely costly, they may also be beneficial or require no intervention (e.g., eutrophic-oligotrophic transition in lakes^16^ and fish recruitment success due to hydro-climate regime shift^46^). Hence, in order to engage in appropriate mitigation actions, it is important to have a plausible estimate of the cost of inaction, which in turn requires us to predict the state of the ecosystem following an impending transition. By making accurate predictions of unobserved dynamical regimes, our approach opens the door to rational decision making and cost-benefit analyses of regime shifts in ecosystems.

## Methods

### Dynamical regimes and bifurcations

Our goal in this study was to predict population dynamics that occurs under unobserved levels of an environmental driver. By doing so, we also provided predictions for the driver levels under which we should observe tipping points. To formalize these ideas, consider a generic discrete-time model that describes the population dynamics of *n* species in an ecosystem:

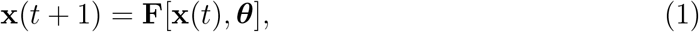

where **x**(*t*) = [*x*_1_(*t*), …, *x*_*n*_(*t*)]^⊤^ is the vector of abundances at time *t*, **F** is a set of *n* functions, and ***θ*** is a set of *m* parameters. Systems modeled as equation (1) can exhibit different types of attractor, including fixed points, cycles, and chaos. Each of these types of attractor may have distinct properties such as high or low abundance for fixed points and long or short periods for cycles. We define a *dynamical regime* as an attractor with a given qualitative property. A *bifurcation* occurs when a small change in one or more *control parameters* in equation (1) causes a change in its dynamical regime— formally, when this parameter change results in a topologically different phase portrait.^14^ We use the terms tipping point and bifurcation interchangeably throughout the text.

Specifically, if *p* ∈ ***θ*** is a control parameter, a bifurcation occurs at a critical value (*p*_*c*_) if we observe distinct dynamical regimes for *p*_*c*_ − *δ* and *p*_*c*_ + *δ*, where *δ* is a small displacement. The number of control parameters that must change to cause a bifurcation is known as the *codimension* of a bifurcation. When working with empirical data, we can leverage knowledge on an environmental driver (e.g., temperature, nutrients, rainfall, fishing pressure) that is known to cause regime shifts and assume that this driver is a proxy for *p*.

In this study, we address the following challenge: having observed time-series data {*x*_*i*_(*t*)} (*t* = 1, …, *T*) for a given species *i* at different dynamical regimes (i.e., at different values of *p*), can we predict regimes at unobserved values of *p* (Fig. 1)? We introduce an approach to solve this challenge by focusing on bifurcations of codimension one. In the next section, we describe our approach based on Empirical Dynamic Modeling (EDM) to predict unseen dynamical regimes from time-series data.

### Approach to predict unseen dynamical regimes

To address the problem described above, we combined Gaussian Process regression with time-delay embedding, an approach known as GP-EDM.^30^ Having observed time series {*x*_*i*_(*t*)} (*t* = 1, …, *T*) for a given species *i* at different values of *p* (Fig. 1a), we would like to learn the following function *F*_*i*_ to be able to predict the dynamics of *x*_*i*_ at unobserved values of *p*:

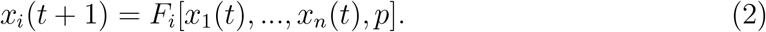

However, if we only have data on species *i*, we propose that we can use time-delay embedding to compensate for unobserved species and instead learn the following function *G*_*i*_ (Fig. 1b):

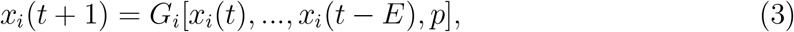

where *E*+1 is the number of lagged versions of *x*_*i*_, also known as the *embedding dimension*. We can then apply an appropriate nonparametric function approximation method, such as Gaussian Process regression, to learn *G*_*i*_ and extrapolate this function to unseen values of *p* (Fig. 1c, d). We provide a detailed description of GP-EDM in *Supplementary Note 1* and we use the R package “GPEDM” to perform our analyses.^47^

Our reasoning for why this approach should work is as follows. Takens’s theorem proves a one-to-one correspondence between an attractor in the native coordinate system (i.e., *x*_1_(*t*), …, *x*_*n*_(*t*)) and one that has been reconstructed using time-delay embedding (i.e., *x*_*i*_(*t*), …, *x*_*i*_(*t* − *E*)).^48^ The bundle embedding theorem^49^ extends time-delay embedding to forced dynamical systems and justifies the inclusion of the control parameter *p* in the reconstruction. Hence, a bifurcation in the native coordinates corresponds to a bifurcation in delay coordinates at the same value of *p*. Therefore, we hypothesize that interpolating between and extrapolating beyond observed values of *p* by learning *G*_*i*_ allows us to reconstruct the bifurcation diagram and to predict the behavior of an ecosystem following a regime shift. Such interpolation and extrapolation should work because, although there is an abrupt change in *x*_*i*_ when crossing a bifurcation, the function *G*_*i*_ should change smoothly when changing *p* (Fig. 1b). Note that if we had data on all species in an ecosystem, then we could use Gaussian Process regression (without time-delay embedding) to learn *F*_*i*_ directly (instead of *G*_*i*_) to predict unseen dynamical regimes.

### Testing approach on model-generated data

We tested our approach on four discrete-time population dynamics models exhibiting different types of bifurcations. These included a single-species model, a two-species competition model with harvesting, a two-species predator-prey model, and a three-species competition model (see next section). For each model, we first determined a range of parameter values, [*p*_*min*_, *p*_*max*_], that produced different dynamical regimes. Then, we selected four distinct values of *p* (*p*_1_, *p*_2_, *p*_3_, and *p*_4_) within [*p*_*min*_, *p*_*max*_] and simulated the dynamics with process noise to generate four time series of length *T* = 50. Note that we iterated each model for 500 time steps and discarded the first 450 points to remove transients. We then stacked these time series together along with additional inputs (i.e., lags of *x*_*i*_ and values of *p*) to create a data matrix to train the GP-EDM model. We standardized all inputs to zero mean and unit standard deviation prior to fitting. We selected the value of *E* of the GP-EDM model based on leave-one-out prediction accuracy, measured as:

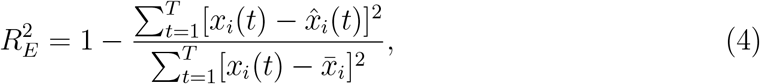

where 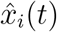 is the GP-EDM prediction for the left-out observation *x*_*i*_(*t*) using a given value of *E* and 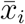 is the mean of *x*_*i*_(*t*) over time. We selected the *E* value that gave the highest prediction accuracy (i.e., max 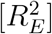 for *E* ∈ {1, …, 9}) across all values of *p*. We also log-transformed *x*_*i*_ if this increased 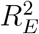.

After we trained the GP-EDM model, we performed predictions for a grid of 200 values of *p* within [*p*_*min*_, *p*_*max*_]. To do so for a given value *p*_*j*_ (*j* = 1, …, 200), we set initial values of *x*_*i*_ and iterated the learned function *G*_*i*_ forward in time while keeping *p* = *p*_*j*_. Specifically, the prediction 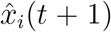 was given by the posterior mean of the GP-EDM model. At every iteration, we used the predicted value as input for the next prediction step. We performed this iteration for 500 time steps and selected the last 100 points as our prediction for the dynamical regime.

We used the Jensen–Shannon divergence averaged over different bin sizes (10 to 15 bins) to measure the accuracy of our predictions. That is, for each value of *p*_*j*_, we had a sample of two distributions of *x*_*i*_, one given by the true time series {*x*_*i*_(*t*)} (*t* = 1, …, *T*) and another given by the predicted time series 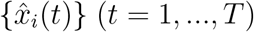, where *T* = 100. Defining *P* (*x*) as the discretized (i.e., binned) probability distribution for {*x*_*i*_(*t*)} and *Q*(*x*) as the discretized probability distribution for 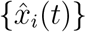, the Jensen–Shannon divergence is measured as:

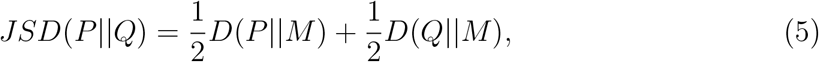

where 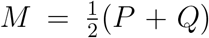 is the mixture distribution of *P* and *Q*, and *D*(*P* ||*M*) is the Kullback–Leibler divergence of *P* from *M* given by:

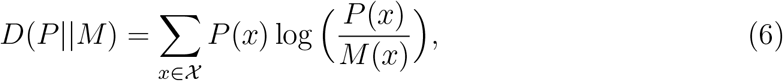

where *χ* is the set of all possible bins and *P* (*x*) denotes the probability (i.e., frequency of points) for bin *x*. Thus, the Jensen–Shannon divergence measures the similarity between two probability distributions (*P* and *Q*), where 0 denotes very similar distributions and 1 denotes very divergent distributions. Note that the true time series in this case is generated from the model without process noise as opposed to the noisy time series of length *T* = 50 that were used to train the GP-EDM model. That is, we reconstructed the bifurcation diagram from noisy data but compared our reconstruction to the true noise-free diagram.

In addition to the analysis described above, we tested whether our approach can be used as an Early Warning Signal (EWS). To do so, we iteratively applied our approach to a time series generated with the two-species competition model under a slowly changing harvest rate (i.e., the control parameter *p*). Specifically, we iterated this model while we increased the harvest rate linearly from 0.42 to 0.48 over 150 time steps. With our parameterization and without process noise (see next section), this model shows a fold bifurcation—that is, a tipping point where the population collapses—at a harvest rate of *p* = 0.459.

Before applying our GP-EDM approach, we performed a simple change-point analysis^50^ to determine the location of the tipping point. We did this because, although the deterministic tipping point occurs at *p* = 0.459, the actual tipping point in the data can occur earlier or later due to process noise. To this end, we split the time series of species *I* into two windows: {*x*_*i*_(1), …, *x*_*i*_(*t*_split_)} and {*x*_*i*_(*t*_split_ + 1), …, *x*_*i*_(*T*)} for 100 values of *t*_split_ within [25, 125]. Then, for each *t*_split_, we computed the sum of squares of window *j* (*SS*_*j*_) using the within-window average, the sum of squares of the entire time series (*SS*_all_) using the global average, and the following ratio 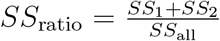. We then selected the *t*_split_ associated with the lowest value of *SS*_ratio_ as the tipping point. The lower the value of *SS*_ratio_, the greater the difference in mean abundance across the two windows.

To apply our GP-EDM approach, we defined 4 time-series windows: (1) *t* = 1, …, 90, (2) *t* = 1, …, 106, (3) *t* = 1, …, 110, and (4) *t* = 1, …, 130. For each window, we trained our GP-EDM model using information on *p* up to the end of the window and then performed predictions for a grid of 300 values of *p* within [0.42, 0.48], exactly as we did in our previous analysis. We also computed the variance of *x*_*i*_(*t*) for successive rolling windows of length 30, which is a well-studied EWS expected to increase before a fold bifurcation. Thus, this analysis allowed us to verify how well we can anticipate, not only an upcoming tipping point, but also the dynamical regime that lies beyond this point.

### Population dynamics models

We used four discrete-time population dynamics models to test our approach. The first model consists of the classic single-species logistic model given by:^51^

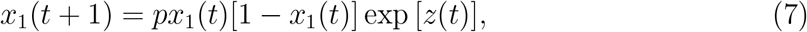

where *p* is the intrinsic growth rate of the population and *z*(*t*) is process noise sampled independently at each time *t* from a normal distribution with mean 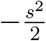 and variance *s*^2^. This noise term is the same for all models (see below) and guarantees that exp [*z*(*t*)] has a log-normal distribution with mean one. For our main set of simulations, we used [*p*_*min*_, *p*_*max*_] = [2.5, 4.0], *p*_1_ = 2.5, *p*_2_ = 2.8, *p*_3_ = 3.25, *p*_4_ = 4.0, and *s* = 0.02.

The second model consists of a two-species competition model with harvesting given by:

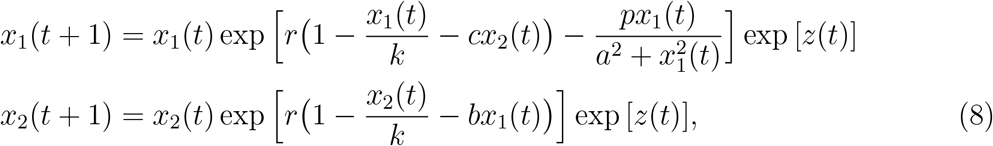

where *r* = 2.7 is the intrinsic growth rate, *k* = 1 is the carrying capacity, *b* = 0.1 and *c* = 0.2 are competition strengths, *p* is the harvest rate, and *a* = 0.1 is the harvest saturation coefficient. For our main set of simulations, we used [*p*_*min*_, *p*_*max*_] = [0.4, 0.5], *p*_1_ = 0.4, *p*_2_ = 0.44, *p*_3_ = 0.47, *p*_4_ = 0.5, and *s* = 0.02.

The third model consists of a two-species predator-prey model given by:^52^

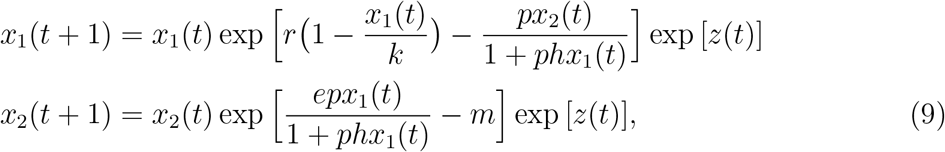

where *r* = 2.5 is the prey intrinsic growth rate, *k* = 4 is the prey carrying capacity, *p* is the predator attack rate, *h* = 0.1 is the predator handling time, *e* = 0.5 is the predator conversion rate, and *m* = 2 is the predator mortality rate. For our main set of simulations, we used [*p*_*min*_, *p*_*max*_] = [2.4, 3.4], *p*_1_ = 2.4, *p*_2_ = 2.6, *p*_3_ = 3.15, *p*_4_ = 3.4, and *s* = 0.02.

The fourth model consists of a three-species competition model given by:^53^

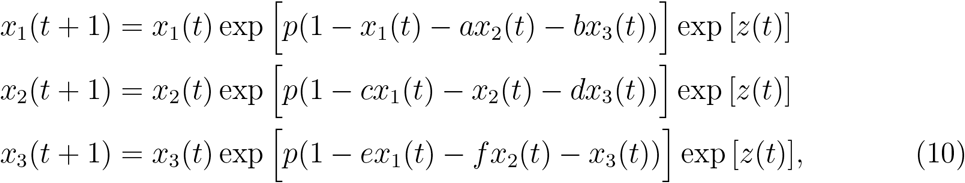

where *p* is the intrinsic growth rate and *a* = 0.5, *b* = 0.5, *c* = 0.1, *d* = 0.2, *e* = 0.2, and *f* = 0.1 are competition strengths. For our main set of simulations, we used [*p*_*min*_, *p*_*max*_] = [1.5, 3.0], *p*_1_ = 1.5, *p*_2_ = 2.1, *p*_3_ = 2.4, *p*_4_ = 3.0, and *s* = 0.02.

### Applying approach to an experimental microbial ecosystem

Our first analysis using empirical data consisted of reconstructing bifurcation diagrams of an experimental microbial ecosystem containing one ciliate predator (*Tetrahymena pyriformis*), a preferred bacterial prey (*Pedobacter* sp.), and a less-preferred bacterial prey (*Brevundimonas* sp.) cultivated in a chemostat.^31^ This data set contains time series for all three species cultivated under five distinct dilution rate treatments. Dilution rates of 0.9/day and 0.75/day resulted in stable fixed points, 0.5/day generated chaos, and 0.45/day resulted in a limit cycle.^31^ We used all experimental replicates, which included one times series for the 0.9/day treatment (*T* = 23), one time series for the 0.75/day treatment (*T* = 28), five time series for the 0.5/day treatment (*T* = 30, *T* = 42, *T* = 42, *T* = 47, and *T* = 52), and two time series for the 0.45/day treatment (*T* = 32 and *T* = 45). Because the only environmental condition changing across treatments is the dilution rate, this data set represents a family of dynamical regimes tuned by a single control parameter, precisely analogous to our simulations shown in Fig. 2.

We applied the same analysis that we performed for model-generated data (see *Testing approach on model-generated data*) independently to each species in this system. For a given species, we first aggregated all replicate time series across the four dilution treatments (i.e., *p* = 0.45, 0.5, 0.75, and 0.9) and included dilution rate (*p*) as an additional input. We removed the 5 initial days of transient dynamics from each replicate to guarantee that each time series covered a single dynamical regime. We then log-transformed abundance as this improved leave-one-out prediction accuracy. Because dilution rate treatments can result in very different mean abundances, we subtracted the mean treatment abundance from each time series. Then, we standardized all inputs and trained the GP-EDM model. We tested *E* ∈ {1, …, 9} and selected the value that maximized leave-one-out prediction accuracy. The selected *E* value was 7 for the predator species, 8 for the preferred prey species, and 8 for the less-preferred prey species. We then extrapolated the GP-EDM model to predict dynamical regimes across a grid of 450 values of *p* within [0.45, 0.9].

### Applying approach to a lake planktonic ecosystem

Our second analysis using empirical data consisted of an EWS analysis to anticipate a potential tipping point from an eutrophic to an oligotrophic regime in a lake planktonic food web. We used a data set of lake planktonic food webs compiled by^16^ and selected Lake Zurich to illustrate our approach, as it contained the longest time series and showed signs of a regime shift driven by a gradual decrease in phosphorus. The planktonic food web from Lake Zurich contained monthly time series for 13 functional groups and 2 environmental variables (phosphate concentration and water temperature) from January 1978 to December 2019 (*T* = 504). Because our EWS analysis with model-generated data was based on a linearly decreasing control parameter, we applied a centered moving average smoother^54^ with a window size of 24 months to remove seasonality and retain the trend in phosphate concentration. We included seasonality in our GP-EDM model through water temperature (see below). Before applying our GP-EDM approach, we performed a change-point analysis^50^ to determine the location of a potential tipping point for each functional group separately (see *Testing approach on model-generated data*). For each functional group, we split its time series into two windows for 300 values of *t*_split_ within [102, 402] and computed *SS*_ratio_ for each *t*_split_. We selected the *t*_split_ associated with the lowest value of *SS*_ratio_ as a potential tipping point.

The above analyses showed that large green algae had the lowest value of *SS*_ratio_ of all functional groups at *t*_split_ = 119 (November, 1987). Thus, we selected the time series window from *t* = 1 (January, 1978) to *t* = 106 (October, 1986) to train our GP-EDM model in order to predict the unseen oligotrophic regime. We first performed leave-one-out predictions for each functional group separately using the trained GP-EDM model to verify which group was more predictable. We used up to 7 lags of functional group abundance (i.e., *x*_*i*_(*t*), …, *x*_*i*_(*t* − 6)), up to 2 lags of water temperature (i.e., *w*(*t*), *w*(*t* − 1)) to account for seasonality, and 1 lag of phosphate concentration (*p*(*t*)) (i.e., control parameter) as inputs in our GP-EDM model. We standardized all inputs before training the GP-EDM model. In addition to showing the lowest value of *SS*_ratio_, large green algae showed the second-best leave-one-out prediction accuracy (*R*^2^ = 0.770). Therefore, we focused on this functional group for subsequent analyses. The best GP-EDM model for large green algae was given by: *x*_*i*_(*t* + 1) = *G*_*i*_[*x*_*i*_(*t*), …, *x*_*i*_(*t* − 4), *w*(*t*), *w*(*t* − 1), *p*(*t*)]. We did not log-transform abundance as this decreased prediction accuracy.

We then followed our EWS analysis with model-generated data (see *Testing approach on model-generated data*) to make predictions for a range of phosphate concentrations using the time series for large green algae. To do so, we defined 4 time-series windows: (1) one year before the potential tipping point (*t* = 1, …, 106), (2) one month before the potential tipping point (*t* = 1, …, 118), (3) four years after the potential tipping point (*t* = 1, …, 166), (4) nine years after the potential tipping point (*t* = 1, …, 226). For each window, we trained the GP-EDM model defined above and then performed predictions for a grid of 300 values of *p* within [0.02, 0.07], which correspond approximately to the minimum and maximum values of phosphate concentration in the data. When iterating predictions forward for a given value of *p*, we fix *p* to that value and set *w* to the average of the corresponding month. By reconstructing the bifurcation diagram for large green algae as more data becomes available, we provide a real-world illustration of our simulations shown in Fig. 3.

## Acknowledgments

We would like to thank Vadim Karatayev, Tanya Rogers, Vasilis Dakos, and Alan Hastings for useful discussions and helpful comments about this work. This work was supported by the Sustainable Oceans National Science Foundation Research Traineeship (grant number #1734999), the Lenfest Oceans Program, and NOAA’s HPCC program.

