## Supplementary Notes and Figures for "Revealing unseen dynamical regimes of ecosystems from population time-series data"

### 1 Gaussian Process regression with time-delay embedding (GP-EDM)

As described in the main text, a key aspect of our approach is to learn the following function $G$:

$$x(t+1) = G[x(t), ..., x(t-E_x), p(t), ..., p(t-E_p)], \tag{S1}$$

where $x(t)$ is the abundance of a given species at time $t$, $p(t)$ is the control parameter at time $t$, $E_x + 1$ is the number of lagged versions of $x$, and $E_p + 1$ is the number of lagged versions of $p$. Note that we dropped the subscript $i$ to denote the $i$-th species in the ecosystem (i.e., $x_i$) to simplify the notation. Also note that in the main text we focused mostly on a single lag for $p$, but the approach can also accommodate multiple lags of $p$, which is particularly useful when the environmental driver changes through time (see *Applying approach to a lake planktonic ecosystem* in the main text). We define $\mathbf{x}(t) = [x(t), ..., x(t-E_x), p(t), ..., p(t-E_p)]^\top$ as the vector containing all lagged versions of $x$ and $p$. If we consider the approximation error $\epsilon(t) \sim \mathcal{N}(0, v)$, we arrive at the following equation:

$$x(t+1) = G[\mathbf{x}(t)] + \epsilon(t), \tag{S2}$$

Note that for a standard implementation of EDM, $\mathbf{x}(t)$ simply includes lags of the state variable we would like to predict (i.e., $\mathbf{x}(t) = [x(t), ..., x(t-E_x)]^\top$). However, in this study, we also include one or more lags of the control parameter. In what follows, we describe how we can use Gaussian Process regression to learn $G$ and extrapolate this function to unseen values of $p$. A similar approach is also described in previous studies such as.[1,2]

A Gaussian Process (GP) is a continuous generalization of the multivariate normal distribution. It is completely specified by a mean function, $m(\mathbf{x}) = \mathbb{E}[G(\mathbf{x})]$, and a covariance function, $C(\mathbf{x}(t), \mathbf{x}(s)) = \mathbb{E}[(G(\mathbf{x}(t)) - m(\mathbf{x}(t)))(G(\mathbf{x}(s)) - m(\mathbf{x}(s)))]$ of the process $G(\mathbf{x})$, where $\mathbf{x}(t)$ and $\mathbf{x}(s)$ represent two different input vectors. A GP is typically written as $G(\mathbf{x}) \sim GP(m(\mathbf{x}), C(\mathbf{x}(t), \mathbf{x}(s)))$.[3] Since we do not have information on the characteristics of $G(\mathbf{x})$ *a priori*, we assumed a constant prior mean function $m(\mathbf{x}) = \mathbf{0}$. In keeping with previous applications of GP-EDM,[1,2] we used a squared-exponential covariance function, given by:

$$C(\mathbf{x}(t), \mathbf{x}(s)) = \tau^2 \exp\Big[-\sum_{j=1}^{D} \phi_j (x(t-j) - x(s-j))^2\Big], \tag{S3}$$

where $\tau^2$ is the prior variance in $G(\mathbf{x})$ and $D$ is the total number of input variables (i.e., dimension of the input vectors). In our case, $D = E_x + E_p + 2$. The $\phi_j$'s are the inverse-length scale parameters, which govern how wiggly the function $G(\mathbf{x})$ is in the direction of the $j$th input. Importantly, if $\phi_j = 0$, then the $j$th input has no effect on the output (i.e., $x(t+1)$).

In keeping with previous applications of GP-EDM, we used minimally informative priors for the hyperparameters and selected values for these by maximizing the marginal posterior distribution after integrating out the unknown function, $G$. We then performed predictions for new values of $\mathbf{x}$ using the posterior distribution for $G$ conditioned on the data and these MAP (maximum *a posteriori*) estimates of the hyperparameters (see[1] for further details and justification).

Our fully specified GP-EDM regression model is given by:

$$\begin{aligned} p(x(t+1) \mid G, \mathbf{x}(t), v) &\sim \mathcal{N}(G(\mathbf{x}(t)), v) \\ p(G \mid \tau^2, \boldsymbol{\phi}) &\sim GP(\mathbf{0}, C) \\ p(\boldsymbol{\phi}, \tau^2, v), \end{aligned} \tag{S4}$$

where $p(\boldsymbol{\phi}, \tau^2, v)$ is the prior specification for the hyperparameters $\boldsymbol{\phi}$, $\tau^2$, and $v$ and the vector $\boldsymbol{\phi} = [\phi_1, ..., \phi_D]^\top$ contains all of inverse-length scale parameters. We used independent, half-normal priors for $\boldsymbol{\phi}$, that is, $p(\phi_j) \sim \frac{2}{\sqrt{2\pi\lambda}} \exp\left[-\phi_j^2/2\lambda\right]$ which has a mode at 0 to encourage irrelevant inputs to drop out of the model. This sparsity-inducing prior is called automatic relevance determination (ARD,[3]) and is analogous to regularization in linear regression. We set $\lambda = \frac{\pi}{2}$ so that the function has approximately one local extremum on average over the range of the data.[1] We set Beta(1.1, 1.1) priors for $\tau^2/Var(x)$ and $v/Var(x)$, which only restricts the total variance in the predicted output to be less than twice the observed variance in the data[1] and facilitates identifiability.

To perform predictions for $x$ using the GP-EDM model, we first determined the hyperparameters that maximize the marginal log likelihood using the time-series data containing $x(t)$ and $p(t)$ for $t = 1, ..., T$, where $T$ is the total number of data points. Defining the vector of hyperparameters as $\boldsymbol{\theta} = [\boldsymbol{\phi}, \tau^2, v]^\top$, the marginal log likelihood is given by:

$$\ln p(\boldsymbol{\theta} \mid \text{data}) = -\frac{1}{2} \ln |C + v\mathbf{I}| - \frac{1}{2} \mathbf{y}^\top (C + v\mathbf{I})^{-1} \mathbf{y} - \ln p(\boldsymbol{\theta}), \tag{S5}$$

where $\mathbf{y}$ is the vector of observed outputs containing $x(t+1)$ for $t = 1, ..., T$ and $\mathbf{I}$ is the $T \times T$ identity matrix. This maximization was done using the R-prop algorithm.[4] Given the estimated hyperparameters, we performed predictions for a set of new inputs $\mathbf{x}_{\text{new}}$

by computing the conditional mean ($m_c$) and covariance ($C_c$) functions evaluated using these new inputs. This was done via the following updating rule:

$$\begin{aligned} m_c(\mathbf{x}_{\text{new}}) &= C(\mathbf{x}_{\text{new}}, \mathbf{X}^\top)(C + v\mathbf{I})^{-1}\mathbf{y} \\ C_c(\mathbf{x}_{\text{new}}, \mathbf{x}_{\text{new}}^\top) &= C(\mathbf{x}_{\text{new}}, \mathbf{x}_{\text{new}}^\top) - C_c(\mathbf{x}_{\text{new}}, \mathbf{X}^\top)(C + v\mathbf{I})^{-1}C(\mathbf{X}, \mathbf{x}_{\text{new}}^\top), \end{aligned} \tag{S6}$$

where $\mathbf{X}$ is the $T \times D$ data matrix containing all input vectors at all time points. We used the R package "GPEDM"[5] to train our GP-EDM models and to perform predictions.

We used equations (S6) to perform leave-one-out predictions and to predict the dynamical regime of $x$ for unseen values of the control parameter $p$. For simplicity, we explain this procedure when using a single lag of $p$. To perform a prediction for a left-out observation $x(t^* + 1)$ at a given time $t^*$, we set $\mathbf{x}_{\text{new}} = [x(t^*), ..., x(t^* - E_x), p(t^*)]^\top$ and used $m_c(\mathbf{x}_{\text{new}})$ from the equations above as our prediction.

To predict the dynamical regime for a given unseen value $\tilde{p}$, we first estimate the hyperparameters using the entire data set for all observed regimes (i.e., all time series of $x$ for all observed values of $p$). Then, we set $\mathbf{x}_{\text{new}} = [x(t^*), ..., x(t^* - E_x), \tilde{p}]^\top$ using the last $E_x + 1$ values of the time series. If we had time series of $x$ for different values of $p$ (e.g., as in Figs. 1 and 2 in the main text), we chose the time series for which the observed $p$ was closest to $\tilde{p}$. Then, we used $m_c(\mathbf{x}_{\text{new}})$ as our prediction for $x(t^* + 1)$. Next, we performed a prediction for $x(t^* + 2)$ using $\mathbf{x}_{\text{new}} = [x(t^* + 1), ..., x(t^* + 1 - E_x), \tilde{p}]^\top$, where $x(t^* + 1)$ is our previous prediction. We repeated this procedure for 500 time steps for our analyses with models and 1000 time steps for our analyses with empirical data. We then used the 100 last predicted abundances as our predicted dynamical regime as explained in the main text.

A final point to note is that when the inverse-length scale parameter for the control parameter is close to zero, the fitted model is insensitive to changes in $p$. This occurred in our initial analyses of the experimental microbial ecosystem (see *Applying approach to an experimental microbial ecosystem* in the main text). As a consequence, the reconstructed bifurcation diagram showed little variation in the dynamics of $x$ as we varied $p$. To circumvent this problem, we fixed the inverse length scale associated with $p$ to 1 for the experimental data and optimized the remaining $\phi_j$ parameters as described above.

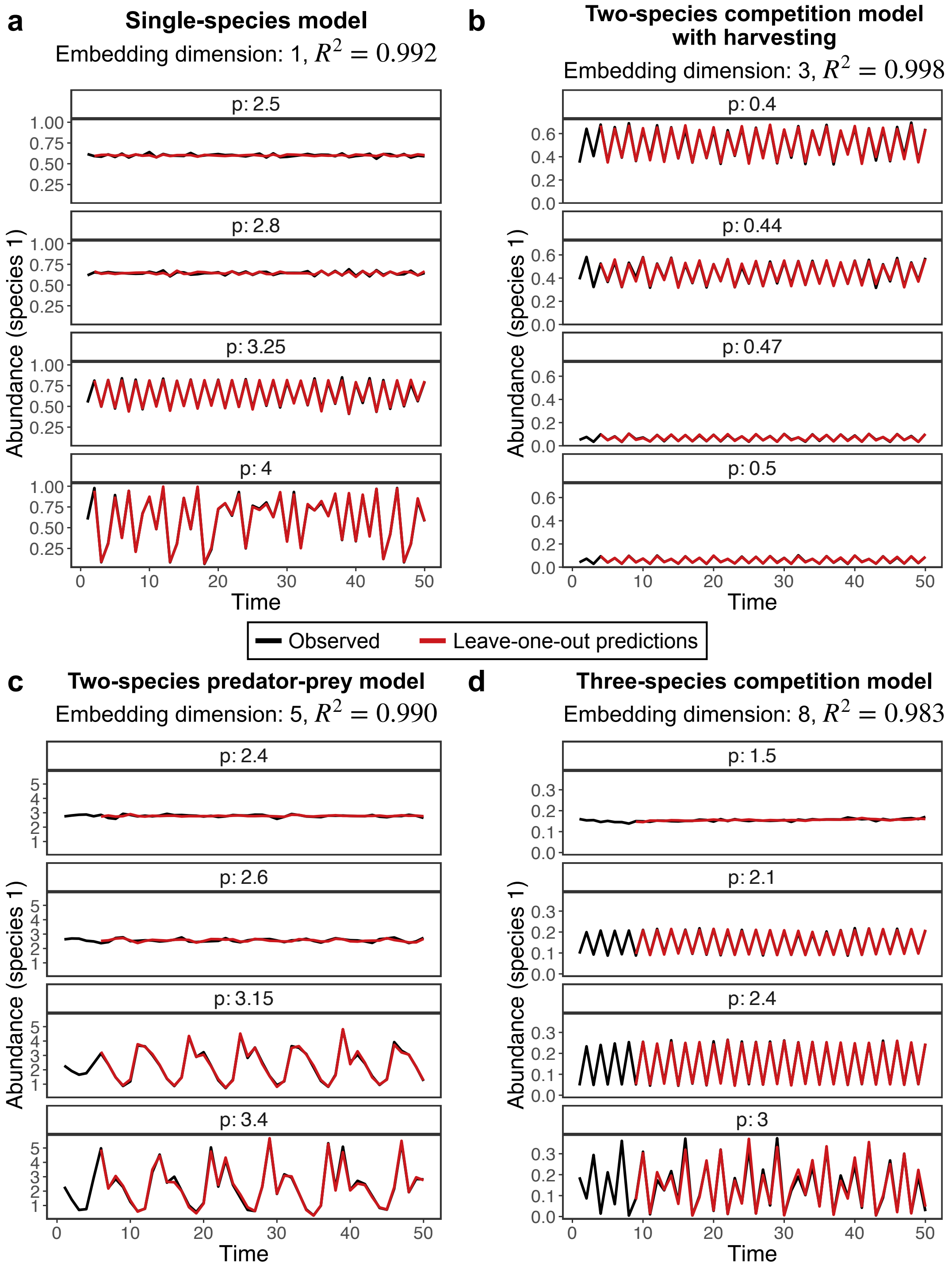


**Fig. S1.** Time series of species 1 ($x_1$, in black) generated from four population dynamics models (equations (5) to (8) in main text) under four different levels of a control parameter ($p$) and the leave-one-out predictions (in red) from GP-EDM. On top of each plot, we list the model used as well as the embedding dimension ($E + 1$) that led to the best prediction accuracy and the associated $R^2$ value.

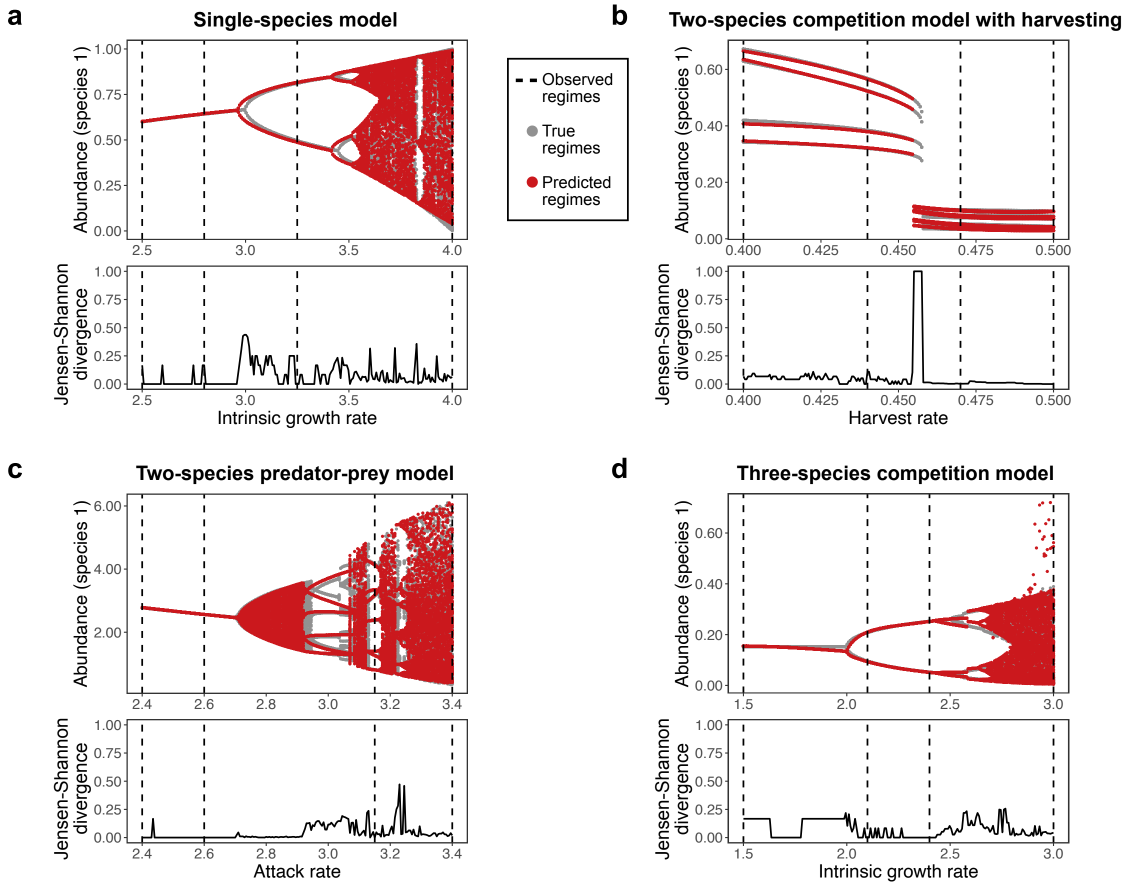


**Fig. S2.** Reconstruction of bifurcation diagrams for all models when we use native instead of delay coordinates with GP-EDM. For example, for the predator-prey model with 2 species, we use Gaussian Process regression to approximate the functions $x_1(t+1) = F_1[x_1(t), x_2(t), p]$ and $x_2(t+1) = F_2[x_1(t), x_2(t), p]$ (i.e., native coordinates) instead of the function $x_1(t+1) = G_1[x_1(t), x_1(t-1), x_1(t-2), x_1(t-3), x_1(t-4), p]$ (i.e., delay coordinates with $E = 4$ as in Fig. 2 in the main text). As expected, the average Jensen-Shannon divergence between true and predicted dynamics was lower with native than with delay coordinates for most models (single-species model, 0.074 vs 0.074; two-species competition model with harvesting, 0.059 vs 0.124; two-species predator-prey model, 0.043 vs 0.042; three-species competition model, 0.077 vs 0.306). Results for delay coordinates are shown in Fig. 2 in the main text.

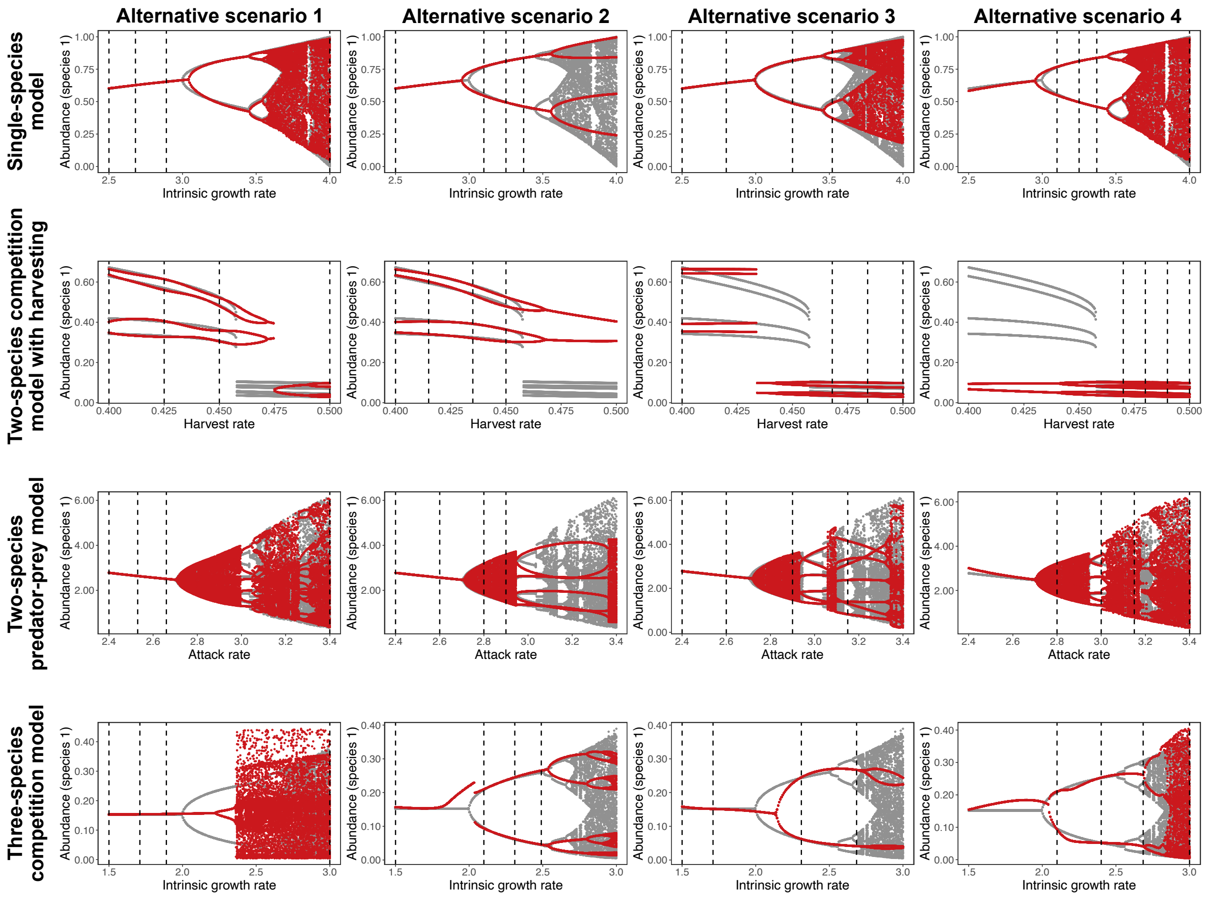


**Fig. S3.** Reconstruction of bifurcation diagrams for all models under four different alternative scenarios of observed dynamical regimes as the training data for GP-EDM. For the single-species model (first row), we use the following scenarios: (1) $p = 2.5, 2.68, 2.89, 4$; (2) $p = 2.5, 3.1, 3.25, 3.37$, (3) $p = 2.5, 2.8, 3.25, 3.52$, and (4) $p = 3.1, 3.25, 3.37, 4$. For the two-species competition model with harvesting (second row), we use the following scenarios: (1) $p = 0.4, 0.425, 0.45, 0.5$; (2) $p = 0.4, 0.415, 0.435, 0.45$, (3) $p = 0.4, 0.468, 0.484, 0.5$, and (4) $p = 0.47, 0.48, 0.49, 0.5$. For the two-species predator-prey model (third row), we use the following scenarios: (1) $p = 2.4, 2.53, 2.66, 3.4$; (2) $p = 2.4, 2.6, 2.8, 2.9$, (3) $p = 2.4, 2.6, 2.9, 3.15$, and (4) $p = 2.8, 3, 3.15, 3.4$. For the three-species competition model (fourth row), we use the following scenarios: (1) $p = 1.5, 1.71, 1.89, 3$; (2) $p = 1.5, 2.1, 2.31, 2.49$, (3) $p = 1.5, 1.71, 2.31, 2.685$, and (4) $p = 2.1, 2.4, 2.685, 3$. In all plots, regimes present in the training data are depicted as vertical dashed lines. The true dynamics are shown in gray and the predicted dynamics in red.

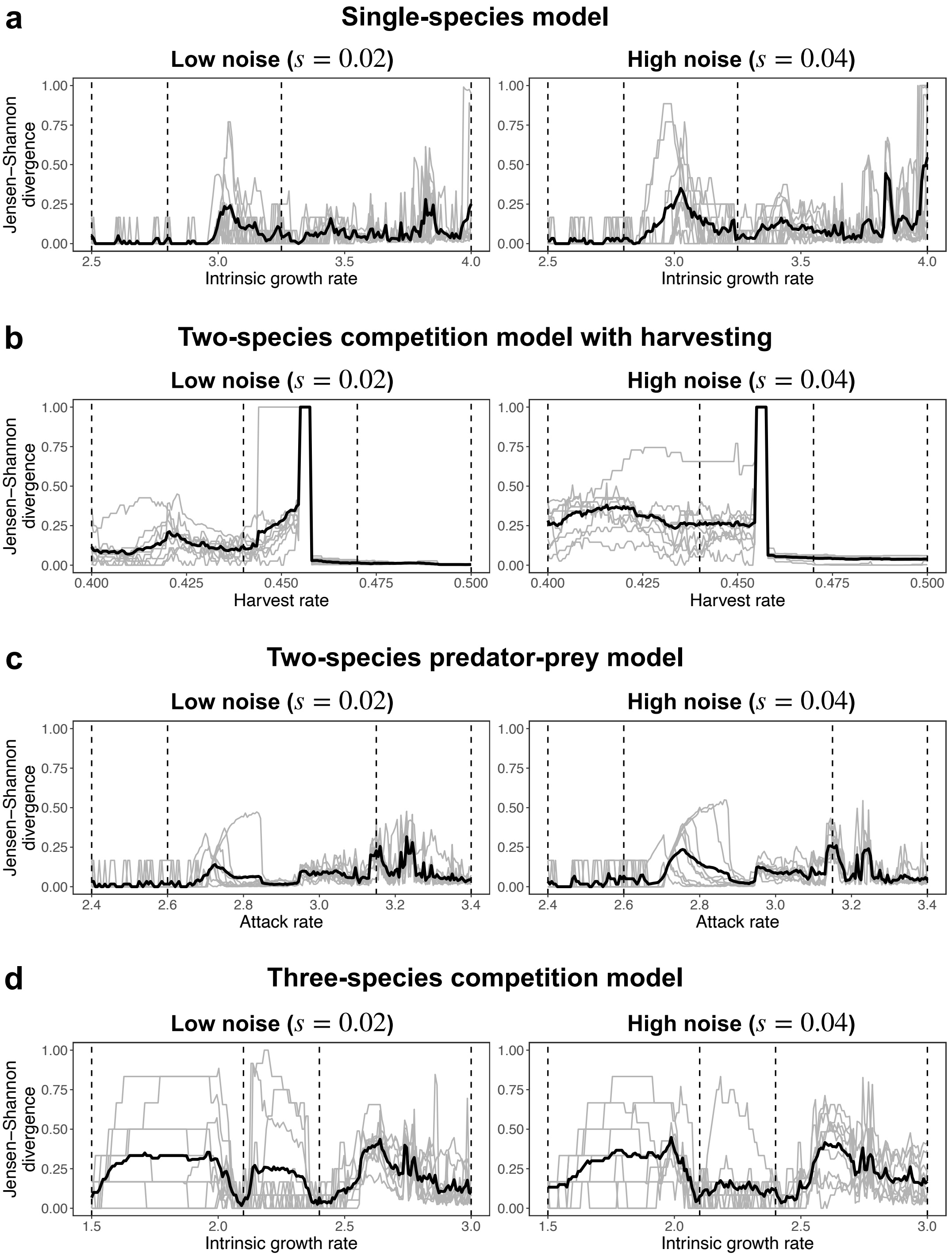


**Fig. S4.** Jensen-Shannon divergence between reconstructed and true bifurcation diagrams for 10 different training data sets and for two levels of process noise for each model. The set up for these analyses is exactly the same as the one described in the main text (see Fig. 2). However, the training data is different because each time we generate the data with process noise, we obtain a different time series. Plots on the left show results for low process noise and plots on the right show results for high process noise. Each plot shows the Jensen-Shannon divergence computed at each value of the control parameter ($p$) for a given training data set (each gray line corresponds to one training data set). The black line corresponds to the average Jensen-Shannon divergence across all training data sets. Values closer to zero represent a closer match between the true and predicted dynamics. In all panels, the vertical dashed lines depict the values of $p$ for which we observed time series to train the GP-EDM model.

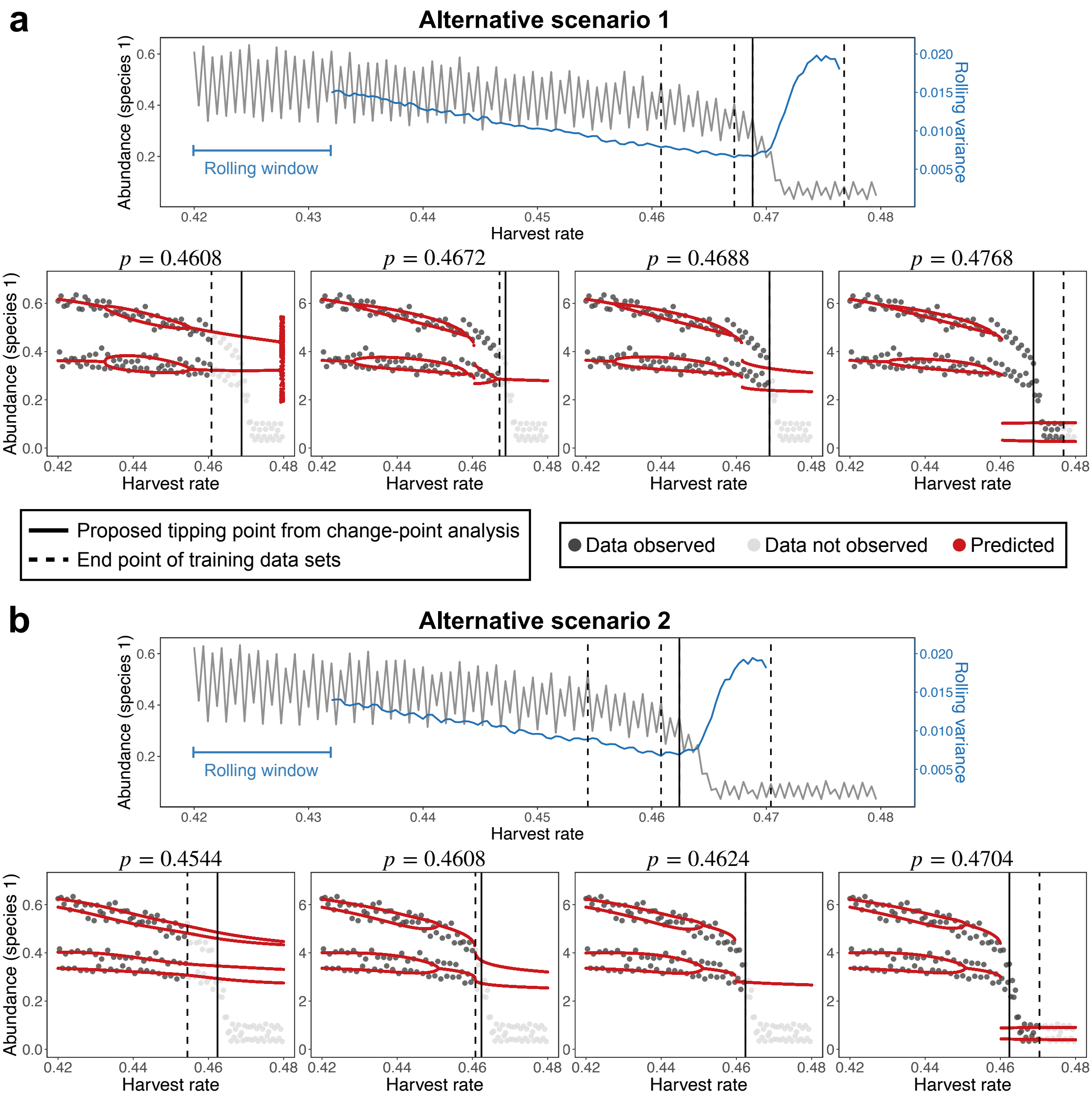


**Fig. S5.** Illustration of GP-EDM approach as an Early Warning Signal using the two-species competition model with harvesting under two alternative training data sets. The set up for these analyses is exactly the same as the one described in the main text (see Fig. 3). However, the training data is different because the location of the tipping point will change each time we generate the time series due to noise. The population (in gray) shows a tipping point at $p = 0.4688$ in **a** and at $p = 0.4624$ in **b** (vertical solid lines) according to a change-point analysis. The vertical dashed lines show the end points of the training data sets used with our GP-EDM model. The rolling variance computed using a window with 30 points is shown in blue. The plots below each time series show the predicted bifurcation diagrams (in red) using the GP-EDM model trained up to the end of each of the four time-series windows. The value of $p$ at the end of the time series window is shown at the top of each plot. Gray points denote population abundance values used to train the GP-EDM model (dark gray) or not yet observed (light gray).

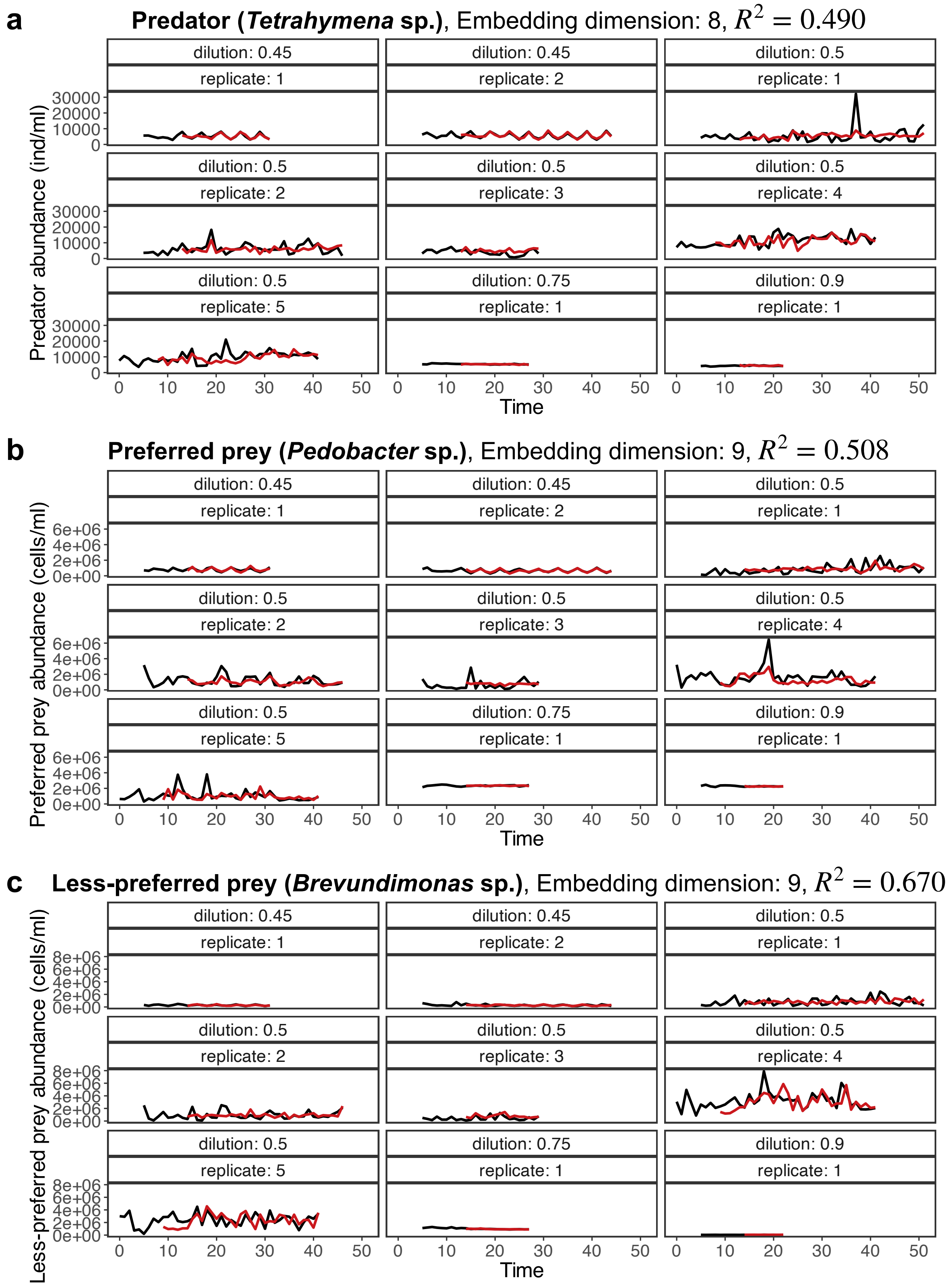


**Fig. S6.** Time series of each species in the experimental microbial ecosystem (in black) under four different levels dilution rate treatments and the leave-one-out predictions (in red) from GP-EDM. For each species, we report the embedding dimension $(E + 1)$ that led to the best prediction accuracy and the associated $R^2$ value.

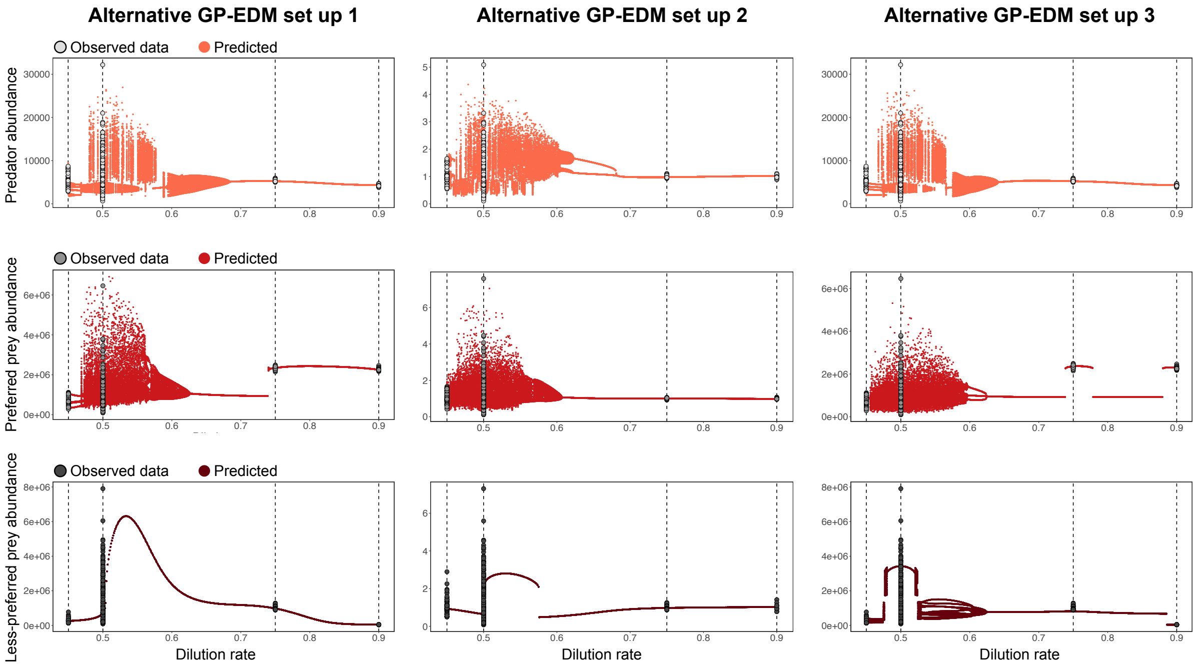


**Fig. S7.** Reconstructed bifurcation diagrams for the 3-species microbial ecosystem using different GP-EDM set ups. Results are qualitatively the same as in Fig. 4 in the main text for the predator and preferred prey species, but not for the less-preferred prey species. The alternative set up 1 is exactly the same as our main text set up (see *Methods* in the main text), but we do not subtract treatment means from each time series prior to standardizing the data. The alternative set up 2 is exactly the same as our main text set up, but we do not fix the inverse-length scale parameter ($\phi_j$) of dilution rate to 1 (see *Gaussian Process regression with time-delay embedding (GP-EDM)*). The alternative set up 3 is exactly the same as our main text set up, but we do not subtract treatment means from each time series prior to standardizing the data and do not fix the inverse-length scale parameter of dilution rate to 1. Note that this last set up is the one used in our analyses with models (i.e., Figs. 2 and 3 in the main text). In all plots, gray points denote the data used to train the GP-EDM model and red points denote predicted dynamical regimes. Vertical dashed lines represent the dilution rates of the experimental treatments (0.45/day, 0.5/day, 0.75/day, and 0.9/day).

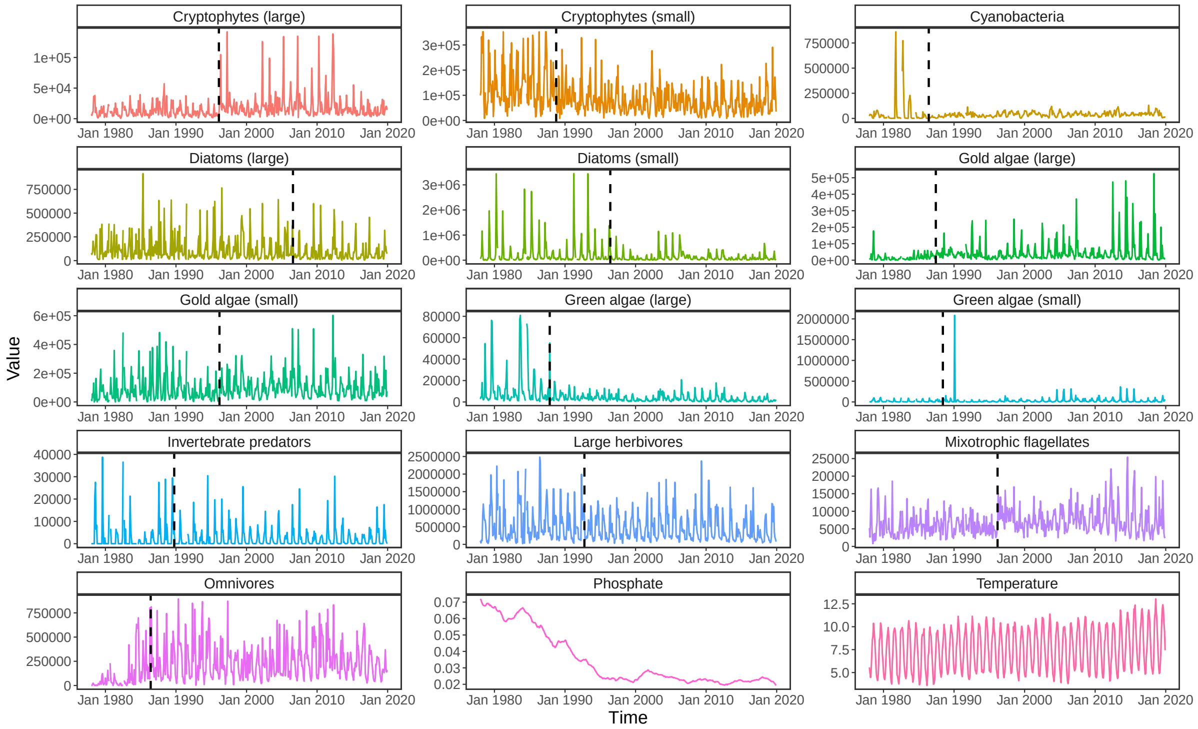


**Fig. S8.** Monthly time series for all 13 plankton functional groups (in ind/l), phosphate concentration (in mg/l), and water temperature (in Celsius) from Lake Zurich between January 1978 and December 2019. As explained in the *Methods* section in the main text, we performed a change-point analysis to determine the location of a potential tipping point for each functional group independently. This potential tipping point is shown for each functional group as a black dashed line and Table S1 gives additional information. For Cryptophytes, small refers to $\leq 700\ \mu m^3$ and large to $> 700\ \mu m^3$. For Diatoms, small refers to $\leq 800\ \mu m^3$ and large to $> 800\ \mu m^3$. For Gold algae, small refers to $\leq 250\ \mu m^3$ and large to $> 250\ \mu m^3$. For Green algae, small refers to $\leq 550\ \mu m^3$ and large to $> 550\ \mu m^3$.

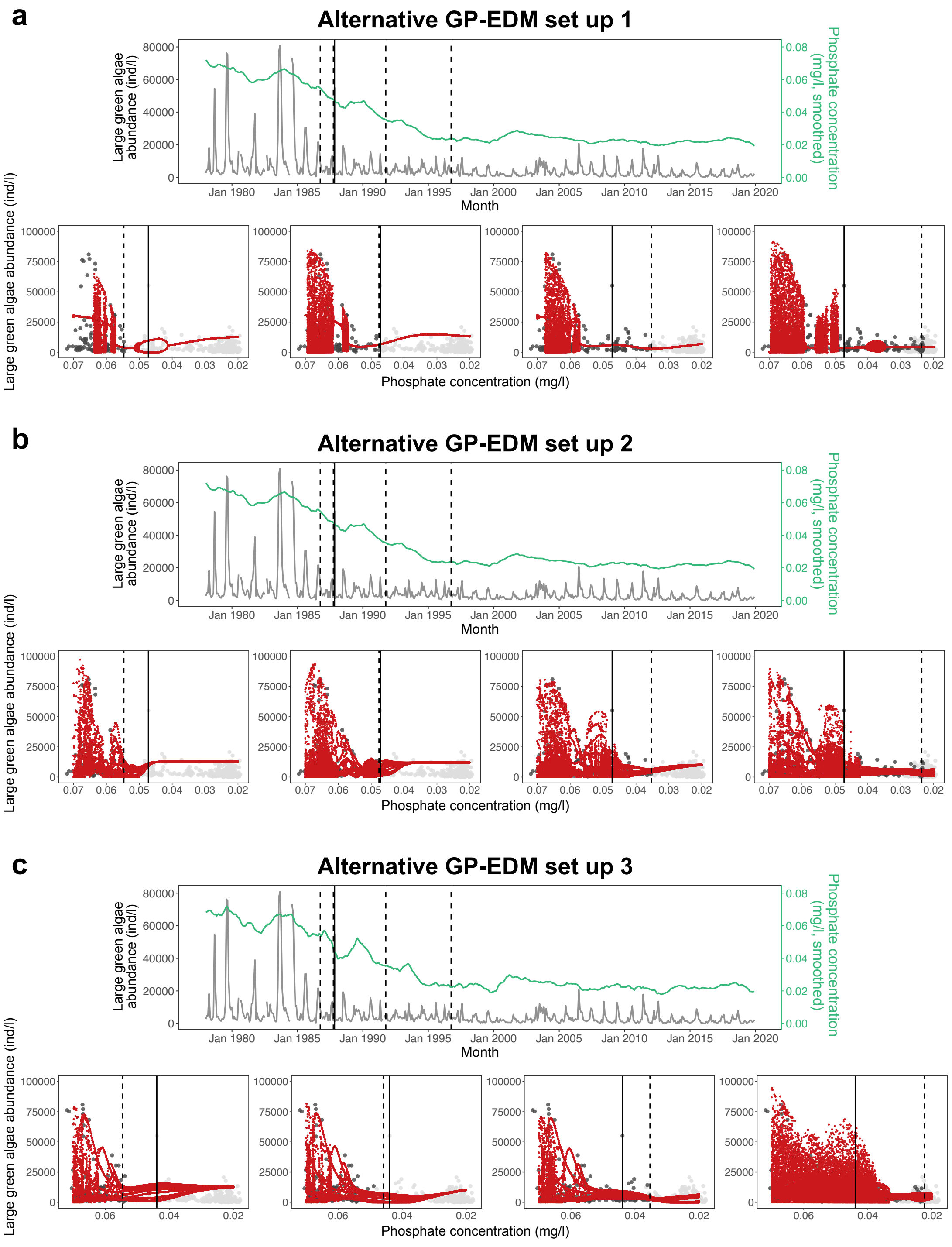


**Fig. S9.** Reconstructed bifurcation diagrams of large green algae in Lake Zurich using different GP-EDM set ups. Results are qualitatively the same as in Fig. 5 in the main text. **a**, Alternative set up 1 is exactly the same as our main text set up, but we do not include temperature as a model input. **b**, Alternative set up 2 is exactly the same as our main text set up, but we fix the inverse-length scale parameter ($\phi_j$) of phosphate to 1. **c**, Alternative set up 3 is exactly the same as our main text set up, but we use a smaller window to smooth the phosphate time series (window with 12 instead of 24 months). In **a**-**c**, the abundance of large green algae is shown in gray and phosphate concentration is shown in green. The potential tipping point is shown as a vertical solid line and vertical dashed lines show the end points of the training data sets. In the bifurcation diagram plots, red points represent our predictions and gray points denote abundance values used to train the GP-EDM model (dark gray) or not yet observed (light gray).

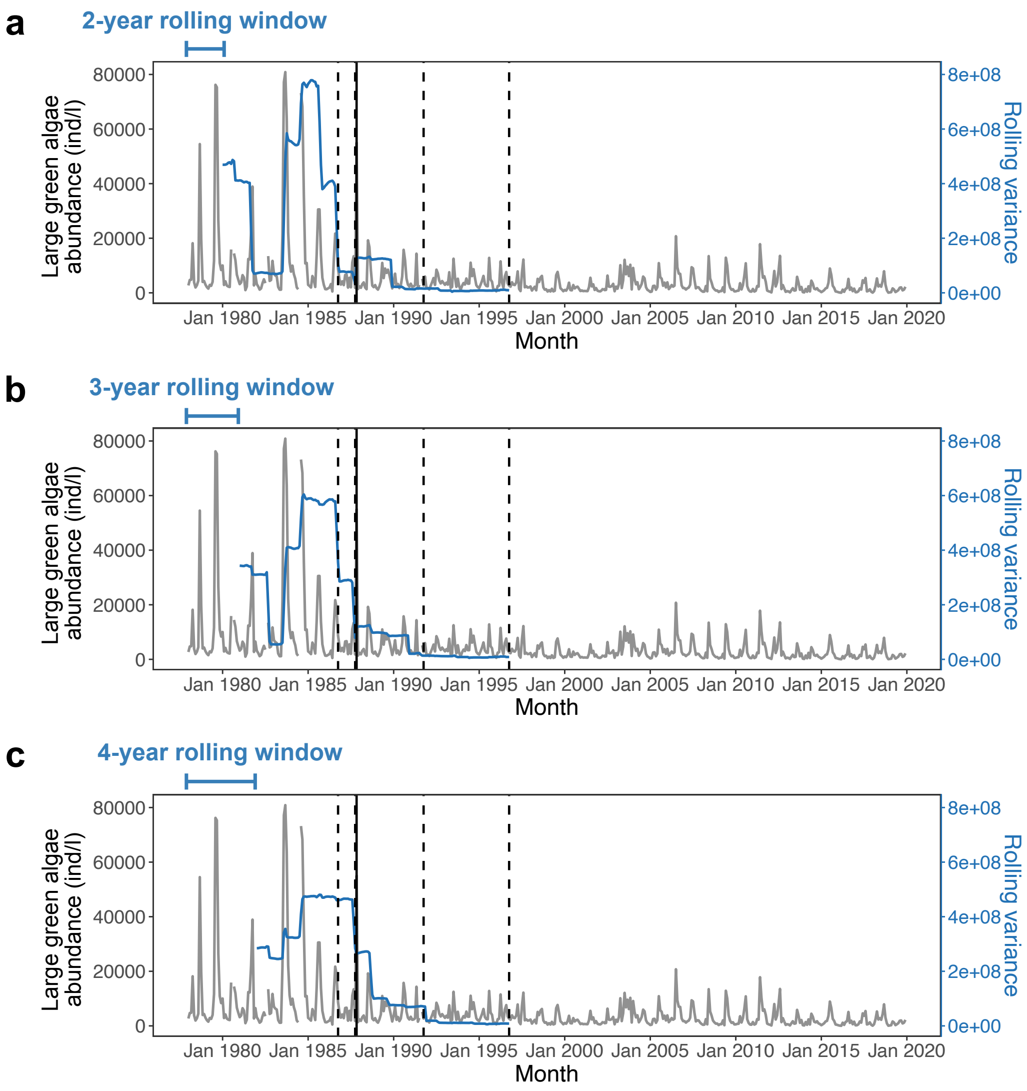


**Fig. S10.** Monthly population time series (in gray) of large green algae and its rolling variance (in blue) in Lake Zurich from January, 1978 to December 2019. The population shows a tipping point on November, 1987 (vertical solid line) according to a change-point analysis. The vertical dashed lines show the end points of the training data sets used with our GP-EDM model. The rolling variance was computed using three different window sizes: 2 years (**a**), 3 years (**b**), and 4 years (**c**).

**Table S1.** Results from the change-point analysis for all plankton functional groups from Lake Zurich (see details in the *Methods* section in the main text). TP time refers to the time point $t$ for the potential tipping point. TP month refers to the month of the potential tipping point. $SS_{\text{ratio}}$ refers to the ratio between $SS_1 + SS_2$ and $SS_{\text{all}}$, where $SS_i$ is the sum of squares of the $i$th time series window and $SS_{\text{all}}$ is the sum of squares of the entire time series. Phosphate refers to the phosphate concentration (in mg/l) at the potential tipping point. We highlight in gray the 3 functional groups with the strongest evidence of a shift in the dynamics, that is, with the lowest values of $SS_{\text{ratio}}$.

| Functional group | TP time | TP month | $SS_{\text{ratio}}$ | Phosphate |
|---|---|---|---|---|
| Cryptophytes (large) | 218 | Feb-96 | 0.940 | 0.024 |
| Cryptophytes (small) | 130 | Oct-88 | 0.855 | 0.043 |
| Cyanobacteria | 102 | Jun-86 | 0.991 | 0.055 |
| Diatoms (large) | 344 | Aug-06 | 0.976 | 0.023 |
| Diatoms (small) | 222 | Jun-96 | 0.961 | 0.023 |
| Gold algae (large) | 114 | Jun-87 | 0.961 | 0.049 |
| Gold algae (small) | 219 | Mar-96 | 0.989 | 0.024 |
| Green algae (large) | 119 | Nov-87 | 0.853 | 0.047 |
| Green algae (small) | 126 | Jun-88 | 0.996 | 0.043 |
| Invertebrate predators | 142 | Oct-89 | 0.995 | 0.046 |
| Large herbivores | 178 | Oct-92 | 0.980 | 0.035 |
| Mixotrophic flagellates | 219 | Mar-96 | 0.931 | 0.024 |
| Omnivores | 102 | Jun-86 | 0.903 | 0.055 |

**Table S2.** Results from the leave-one-out prediction analyses for all plankton functional groups from Lake Zurich (see details in the *Methods* section in the main text). For each functional group, we show the details of the selected GP-EDM model, that is, the model with the highest $R^2$ value. Embedding dimension refers to the total number of lagged inputs in the GP-EDM model (lags of abundance + lags of temperature + lags of phosphate). $\boldsymbol{\phi}$ refers to the set of inverse length scale parameters ($\phi_j$) for all model inputs. $R^2$ is the coefficient of determination computed from the GP-EDM predictions. We highlight in gray the 3 most predictable functional groups, that is, with the highest values of $R^2$.

| Functional group | Embedding dimension | $\boldsymbol{\phi}$ | $R^2$ |
|---|---|---|---|
| Cryptophytes (large) | $6+2+1=9$ | 0, 0.022, 0, 0, 0, 0, 0.388, 0.402, 0 | 0.405 |
| Cryptophytes (small) | $6+2+1=9$ | 0.397, 0, 0, 0.075, 0.122, 0.572, 0, 0.331, 0.04 | 0.394 |
| Cyanobacteria | $1+1+1=3$ | 0.405, 0.179, 0 | 0.696 |
| Diatoms (large) | $4+2+1=7$ | 0, 0, 0.417, 0, 0.236, 1.158, 0 | 0.186 |
| Diatoms (small) | $1+2+1=4$ | 2.392, 1.556, 0, 0.055 | 0.492 |
| Gold algae (large) | $5+1+1=7$ | 0.269, 0, 0.18, 0.437, 0, 0.424, 0 | 0.174 |
| Gold algae (small) | $7+1+1=9$ | 0.453, 0, 0, 0, 0.068, 0.072, 0.006, 0.609, 0.015 | 0.271 |
| Green algae (large) | $4+2+1=7$ | 0.713, 0.164, 0.196, 0.621, 0, 0.352, 0.066 | 0.77 |
| Green algae (small) | $6+2+1=9$ | 0.289, 0.077, 0, 0.176, 0, 0.23, 0.165, 1.011, 0.022 | 0.801 |
| Invertebrate predators | $6+2+1=9$ | 1.368, 0.005, 0, 0.037, 0, 0, 0.471, 1.494, 0.644 | 0.325 |
| Large herbivores | $6+2+1=9$ | 0.015, 0, 0, 0, 0, 0.017, 0.337, 0.111, 0.424 | 0.526 |
| Mixotrophic flagellates | $6+2+1=9$ | 0.157, 0, 0, 0, 0.841, 0, 0.222, 0.11, 0 | 0.46 |
| Omnivores | $4+2+1=7$ | 0.042, 0.219, 1.173, 1.145, 0, 0.326, 0 | 0.399 |
